# Perivascular and peribronchiolar granuloma-associated lymphoid tissue and B-cell gene expression pathways identify asymptomatic *Mycobacterium tuberculosis* lung infection in Diversity Outbred mice

**DOI:** 10.1101/2023.07.27.550843

**Authors:** Deniz Koyuncu, Thomas Tavolara, Daniel M. Gatti, Adam C. Gower, Melanie L. Ginese, Igor Kramnik, Bülent Yener, Muhammad Khalid Khan Niazi, Metin Gurcan, Anas Alsharaydeh, Gillian Beamer

## Abstract

Humans are highly genetically diverse, and most are resistant to *Mycobacterium tuberculosis.* However, lung tissue from genetically resistant humans is not readily available to identify potential mechanisms of resistance. To address this, we model *M. tuberculosis* infection in Diversity Outbred mice. Like humans, Diversity Outbred mice also exhibit genetically determined susceptibility to *M. tuberculosis* infection: Progressors who succumb within 60 days of a low dose aerosol infection due to acute necrotizing granulomas, and Controllers who maintain asymptomatic infection for at least 60 days, and then develop chronic pulmonary TB with occasional necrosis and cavitation, over months to greater than 1 year. Here, we identified specific regions of granuloma-associated lymphoid tissue (GrALT) and B-cell gene expression pathways as key features of asymptomatic lung infection using cytokine, antibody, granuloma image, and gene expression datasets. Cytokines and anti-*M. tuberculosis* cell wall antibodies discriminated acute vs chronic pulmonary TB but not asymptomatic lung infection. To find unique features of asymptomatic lung infection, we trained a weakly supervised, deep-learning neural network on lung histology images. The neural network accurately produced an interpretable imaging biomarker: perivascular and bronchiolar lymphocytic cuffs, a type of GrALT. We expected CD4 T cell genes would be highly expressed in asymptomatic lung infection. However, the significantly different, highly expressed genes in lungs of asymptomatically infected Diversity Outbred mice corresponded to B-cell activation, proliferation, and antigen-receptor signaling, including *Fcrl1, Cd79, Pax5, Cr2,* and *Ms4a1*. Overall, our results suggest that genetically controlled B-cell responses are important for establishing asymptomatic *M. tuberculosis* lung infection.

## INTRODUCTION

Tuberculosis (TB) is a globally important disease. Over 2 billion people harbor infection with *Mycobacterium tuberculosis*, resulting in 9-10 million new TB cases and 1.6 million deaths [1]. Humans respond variably to *M. tuberculosis* infection, ranging from extreme susceptibility to extreme resistance. Only 5-10% of infected adult humans ever develop pulmonary disease: Very rarely developing acute fulminant TB; and most commonly developing post-primary TB, a chronic and destructive lung disease that can wax and wane [2-5]. Most (90-95%) humans are highly resistant to *M. tuberculosis* and clear bacilli by innate immunity [6], or restrict bacillary growth by adaptive immunity and maintain asymptomatic infection or latent TB infection [7]. Severe immune deficiency, malnutrition, vitamin D deficiency, diabetes, extreme age, smoking, illicit drug use, and co-infections are known risk factors for pulmonary TB; however, their presence does not fully explain susceptibility, and their absence does not fully explain resistance [8-11].

Additional host factors, such as genetically controlled responses contribute to resistance and susceptibility to *M. tuberculosis* [12, 13]. Large effects due to host genetic background have been shown in immune-competent mouse models by using panels of Collaborative Cross inbred mouse strains [14, 15] and by using the Diversity Outbred mouse population [16-18]. The Diversity Outbred population’s genetic diversity, heterozygosity, and phenotypic ranges are comparable to humans, providing a valuable small animal model to study phenotypic heterogeneity and investigate the genetic basis of complex traits [19]. We and others have shown that aerosolized *M. tuberculosis* causes severe, acute necrosuppurative inflammation that does not restrict *M. tuberculosis* growth in ∼30% of Diversity Outbred mice and these Progressors succumb to acute pulmonary TB within 8 weeks [16, 18, 20]. The remaining ∼70% of Diversity Outbred mice survive longer, have significantly lower levels of acute inflammation, and develop non-necrotizing lung granulomas that better restrict *M. tuberculosis* [16, 18, 21, 22].

Our prior work used multiple approaches to study Progressors, including granuloma analysis, transcriptional profiling, and statistical and machine learning methods to find and validate translational diagnostic biomarkers [18, 23]. We developed and validated a weakly supervised, attention-based, multiple instance learning model that diagnosed Progressors with high accuracy (91.50 ± 4.68%) from digital granuloma images. Further, *post hoc* visual examination by board-certified pathologists confirmed that the “imaging biomarker” of Progressors corresponded to granuloma regions of pyknotic nuclear debris [24]. Transcriptional analyses identified a unique, highly expressed, inflammatory lung signature which subsequently led to discovery and validation of diagnostic biomarkers for pulmonary TB, one of which (serum CXCL1) met the World Health Organization’s diagnostic criteria for a triage test in human sera [23].

Here, we extend that work by showing the culmination of large survival studies lasting more than six hundred days, and novel approaches to uncover features of resistance to *M. tuberculosis* in Diversity Outbred mice. We distinguished two types of resistance: (i) Lung resistance in *asymptomatically* infected Diversity Outbred mice who had no clinical signs of disease at the experimental endpoint on or before 60 days post-infection; and (ii) Host resistance in *Controllers* where infection was allowed to progress, and Controller survived months to years. Controllers are genetically resistant hosts, and when euthanized due to morbidity, their lungs model end stage, chronic, pulmonary TB.

## METHODS

### Ethics Statement

Tufts University’s Institutional Animal Care and Use Committee (IACUC) approved this work under protocols G2012-53; G2015-33; G2018-33; and G2020-121. Tufts University’s Institutional Biosafety Committee (IBC) approved this work under registrations: GRIA04; GRIA10; GRIA17, and 2020-G61.

### Mice

Four-to-five-week old female Diversity Outbred mice (n=1009) from generations 15, 16, 21, 22, 34, 35, 37, and 42 were purchased from The Jackson Laboratory (Bar Harbor, ME) and group housed (n=5-7 mice per cage) on Innovive (San Diego, CA) or Allentown Inc (Allentown, NJ) ventilated, HEPA-filtered racks in the New England Regional Biosafety Laboratory (Tufts University, Cummings School of Veterinary Medicine, North Grafton, MA). The light cycle was 12 hours of light; 12 hours of dark. Disposable caging was purchased sterile and re-usable caging was autoclaved prior to use. All cages were lined with sterile corn-cob bedding and sterile paper nestlets (Scotts Pharma Solutions, Marlborough, MA). Cages were changed at least every other week. Mice were provided with sterile mouse chow (Envigo, Indianapolis, IA) and sterile, acidified water *ad libidum*.

### *M. tuberculosis* aerosol infection

At eight-to-ten-weeks old, mice were infected with aerosolized *M. tuberculosis* strain Erdman using a custom-built CH Technologies system [18, 23, 25]. Twenty-four hours after each aerosol run, 4-12 mice were euthanized by carbon dioxide, and the entire lungs homogenized in 5mL sterile phosphate buffered saline, and all homogenate plated onto OADC-supplemented 7H11 agar. After 3 weeks at 37°C, *M. tuberculosis* colony forming units were counted to determine the retained lung dose [18, 23, 25]. Mice were infected with ∼100 bacilli in the first two experiments, and ∼25 bacilli in the subsequent eight experiments.

### Clinical monitoring and survival

Mice were observed daily for routine health monitoring. Mice were weighed before *M. tuberculosis* aerosol infection, and at least once per week afterward until consecutive weight loss was noted, and then mice were weighed up to daily. Criteria requiring euthanasia were any one of the following: severe weakness/lethargy; respiratory difficulty; or body condition score < 2 [26]. We confirmed morbidity was due to pulmonary TB by finding: (i) Large nodular, or severe diffuse lung lesions; (ii) Histopathological confirmation of neutrophilic, lymphoplasmacytic, histiocytic, or granulomatous lung infiltrates; (iii) Cultivable *M. tuberculosis* bacilli from lung tissue; and (iv) Absence of other disease based on necropsy findings. Since IACUC protocols prohibited death as an endpoint, the day a mouse was euthanized due pulmonary TB was used as a proxy of survival. Mice with morbidity not attributable to pulmonary TB were excluded from subsequent analyses.

### Quantification of *M. tuberculosis* lung burden

Immediately after euthanasia, two or three lung lobes were removed from each mouse and homogenized in 1mL sterile phosphate buffered saline per lobe, serially diluted, plated onto OADC-supplemented 7H11 agar, and incubated at 37°C. After 3-4 weeks, *M. tuberculosis* colonies were counted, and *M. tuberculosis* burden in the lungs calculated [18, 23, 25].

### Routine histology

One or two lung lobes from each Diversity Outbred mouse were inflated and fixed in 10% neutral buffered formalin, processed, embedded in paraffin, sectioned at 5µm, and stained with carbol fuschin for Acid-Fast bacteria followed by counterstaining with hematoxylin and eosin (H&E) at the Cummings School of Veterinary Medicine’s Comparative Genomics and Pathology Shared Resource (North Grafton, MA). Stained tissue sections on glass slides were digitally scanned by Aperio ScanScope or AT2 scanners at 0.23 microns per pixel at Vanderbilt University Medical Center’s Digital Histology Shared Resource (Nashville, TN).

### Computer-aided image analysis using attention-based multiple instance learning

Lung tissue section images from 129 *M. tuberculosis-*infected Diversity Outbred mice were used for training and 98 different images were used as a hold-out test set. Of the 129 training images, 66 were from Asymptomatic mice and 63 images were from Controllers; of the 98 test set images 10 were Asymptomatic mice and 88 were controllers. Following image preprocessing (see Supplementary Methods), we used an attention-based multiple instance learning method [27]. The attention weights corresponded to regions of the images that contributed to classification and yielded an interpretable model [24]. Briefly, we assigned lung tissue images based on the two host classes and trained an end-to-end deep learning model to predict the image-level class. The model consisted of two parts: a feature extractor followed by the attention mechanism, depicted in Supplementary Figure 5. Optimization used the Adam optimizer with the following parameters: β1 = 0.9 and β2 = 0.999, the learning rate of 0.0001, weight decay of 0.0005, and over 100 epochs. For each fold, 30 Diversity Outbred mice were randomly sampled for the validation set, and the rest of the cases were used for the training of the model, known as Monte Carlo cross-validation [28]. To account for imbalanced training sets, the training procedure was modified to randomly select images from Controller or Asymptomatic mice with equal likelihood and then to randomly select a mouse from the selected category during every training iteration. Negative log-likelihood was used as a cost function. The code used in the analysis will be made publicly available upon publication.

### Gene expression in lung tissue by microarray analysis

One lung lobe from n=117 Diversity Outbred mice was homogenized in TRIzol™, stored at -80°C, and RNA was extracted using PureLink RNA Mini Kits (Life Technologies, Carlsbad, CA). The Boston University Microarray and Sequencing Resource Core Facility (Boston, MA) confirmed RNA quality and quantity and prepared and hybridized material to Mouse Gene 2.0 ST microarrays. Raw CEL files were normalized to produce gene-level expression values using the implementation of the Robust Multiarray Average (RMA) in the affy R package (version 1.62.0) and an Entrez Gene-specific probeset mapping (17.0.0) from the Molecular and Behavioral Neuroscience Institute (Brainarray) at the University of Michigan [29]. All microarray data processing was performed using the R environment for statistical computing (version 3.6.0).

### Quantification of lung cytokines, chemokines, and anti-*M. tuberculosis* antibodies

After plating for *M. tuberculosis* colony forming units, remaining lung homogenates from all mice were aliquoted and stored at -80°C. Homogenates were thawed overnight at 4°C, serially diluted and assayed by sandwich ELISA for cytokines and chemokines (CXCL5, CXCL2, CXCL1, TNF, MMP8, S100A8, IFN-γ, IL12p40, IL-12p70, IL-10, and VEGF using antibody pairs and standards from R&D Systems (Minneapolis, MN), Invitrogen (Carlsbad, CA), eBioscience (San Diego, CA), or BD Biosciences (San Jose, CA, USA), per kit instructions. A subset of these ELISA results was reported previously [23]. The amount of immunoglobulin G (IgG) bound by *M. tuberculosis* cell wall fraction; *M. tuberculosis* culture filtrate proteins; *M. tuberculosis* Antigen 85 Complex; and *M. tuberculosis* ESAT-6:CFP-10 complex was quantified using in-house optimized ELISAs [23, 30]. Briefly, high-binding immunoassay plates (Corning Costar #9018) were coated overnight in 100µl per well of 1-5 µg/mL *M. tuberculosis* cell wall fraction (NR-14828); *M. tuberculosis* culture filtrate proteins (NR-14825); purified native *M. tuberculosis* Antigen 85 complex (NR-14855); or recombinant purified *M. tuberculosis* Early Secreted Antigenic Target (ESAT)-6 (NR-49424) and *M. tuberculosis* Culture Filtrate Protein (CFP)-10 (NR-49425) obtained through BEI Resources, NIAID, NIH. The following day, plates were blocked, lung homogenates serially diluted in 1% Bovine Serum Albumin (BSA), and incubated overnight at 4°C. After multiple washes, goat anti-mouse IgG-HRP (Rockland Immunochemicals #610-13) was diluted per manufacturer’s instructions, incubated, washed multiple times, developed with TMB (ThermoFisher or R&D Systems), stopped using 0.25M HCl, and read at 450nm using a BioTek Plate reader. Concentration of bound IgG was computed based on standard curves and 4-parameter logistic regression models. Data will be made available upon publication.

### Classification using cytokines, chemokines, and anti-*M. tuberculosis* IgG

#### Data

For the cytokines, chemokines, and IgG antibody-based classification, we filtered for the mice with complete measurements for the twelve cytokines and antibodies: CXCL5, CXCL2, CXCL1, TNF, IFNg, IL-12, IL-10, MMP8, S100A8, VEGF, anti-*M. tuberculosis* CW, and anti-*M. tuberculosis* CFP). This filtering yielded n=30 Asymptomatic; n=48 Controllers, and n=38 Progressor mice from two independent experimental infections.

#### Classification methods

To discriminate between acute pulmonary TB in Progressors, chronic pulmonary TB in Controllers, and asymptomatic lung infection, we first used a linear classifier, L1 regularized logistic regression. The regularization term promotes sparse coefficients [31] and *λ* is selected through grid search among {0, 10^-2^, 10^-1.95^,…, 10^1.95^, 10^2^}. We have used the scikit-learn implementation [32] of the logistic regression model. To mitigate the unbalanced classes, sample reweighing with the “balanced” option of the scikit-learn library is used. We defined feature importance of a biomarker as the ratio of its (absolute) effect size (defined below) to the sum of all (absolute) effect sizes. More formally, let *β*_’_ denote the effect size of the j’th biomarker, its feature importance is given by 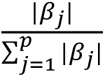 where p is the total number of biomarkers in that panel (i.e., 10 or 12). As an alternative method we also tested a non-linear classifier, XGBoost [33]. We have used the python interface of the XGBoost and grid searched the two parameters: “learning_rate” ([1e-3,1e-2]) and “n_estimators” ([3,5,100]). We fixed the parameter “max_depth” to 3 and used the default values for the remaining. During training, we again used sample reweighing. We reported the performance corresponding to the classifier’s best hyper-parameter using 30-fold-cross-validation. Further details are discussed in the Supplementary Methods. The code used in the analysis will be made publicly available upon publication.

### Statistical analyses and other performance metrics

#### Data from *M. tuberculosis* infected mice

Survival, weight loss, *M. tuberculosis* lung burden, and ELISA data were analyzed and graphed in GraphPad Prism 8.4.2 with significance set at p < 0.05 and adjusted for multiple comparisons. Survival curves were analyzed using Log-rank (Mantel-Cox) test. Body weight data, lung *M. tuberculosis* burden, and lung cytokines, chemokines, and antibodies were analyzed for normal or lognormal distributions prior to one-way ANOVA by Kruskal-Wallis with Dunn’s post-test or Brown-Forsythe and Welch’s with Dunnett’s T3 post-test as indicated in the Figure legends.

#### Imaging biomarkers

Model performance was evaluated using overall sensitivity, specificity, and Area Under the Receiver Operating Characteristic Curve (AUC) of a ten-fold Monte-Carlo cross-validation. 95% confidence intervals for each statistic were computed using bootstrapped samples of predictions (equal to the number of observations) with replacement (n = 1000). 97.5th and 2.5th percentiles were taken as bounds for confidence intervals.

#### Gene expression analyses

For each gene, AUC analysis was performed using R package pROC (v1.18) [34]. Corresponding p-values to the AUC, are calculated using one sided Mann–Whitney U-statistic [35] with Python package statsmodels (v0.13.2) [36]. In the Supplementary Methods, we compared the statistically significant genes resulting from Mann–Whitney U-statistic with a parametric alternative, Welch’s t-test. For each classification problem, the directionality of the test was selected such that the gene expression values were statistically higher in the class with longer survival under the alternative hypothesis. Benjamini-Hochberg correction was applied separately to each classification problem to control the false discovery rate at 0.05. Clustermap is drawn using the python package seaborn (v0.11.2) [37] with clustering metric “correlation”. Enrichr (https://amp.pharm.mssm.edu/Enrichr) was used to identify Gene Ontology (GO) biological processes (version 2021) that were significantly overrepresented (adjusted *p* < 0.05) within an input set of official mouse gene symbols. A subset of microarray data and analyses from Progressor mice were published elsewhere [23, 38], deposited in Gene Expression Omnibus (GEO) and assigned Series ID GSE179417. Deposition of the gene expression dataset described in this study in the Gene Expression Omnibus (GEO) is pending.

## RESULTS

### Survival, *M. tuberculosis* lung burden, and inflammatory biomarkers in infected mice

A low dose of aerosolized *M. tuberculosis* (20 ± 12 bacilli) results in early morbidity and mortality in approximately one-third of Diversity Outbred mice, and the supersusceptible Progressor mice succumb to acute necrosuppurative pulmonary TB with high bacterial burden within 60 days of infection. This phenotype is reproducible across sexes, institutions, aerosol infection methods, and strains of *M. tuberculosis* and is not due to immune deficiency [16, 18, 20, 23]. *M. tuberculosis* infection significantly reduces survival of Diversity Outbred mice compared to identically housed, age- and sex-matched non-infected Diversity Outbred controls; and survival is significantly different than *M. tuberculosis-*infected C57BL/6J inbred mice, with approximately 25% of Diversity Outbred mice surviving longer than the median survival of C57BL/6J (Figure 1A). The ∼70% of Diversity Outbred mice that are more resistant to *M. tuberculosis* (i.e., Controllers) survive longer than Progressors (Figure 1B). Table 1 summarizes susceptibility and resistance to *M. tuberculosis* in humans [39-42] that may be comparable to Diversity Outbred mice.

**Figure 1.**
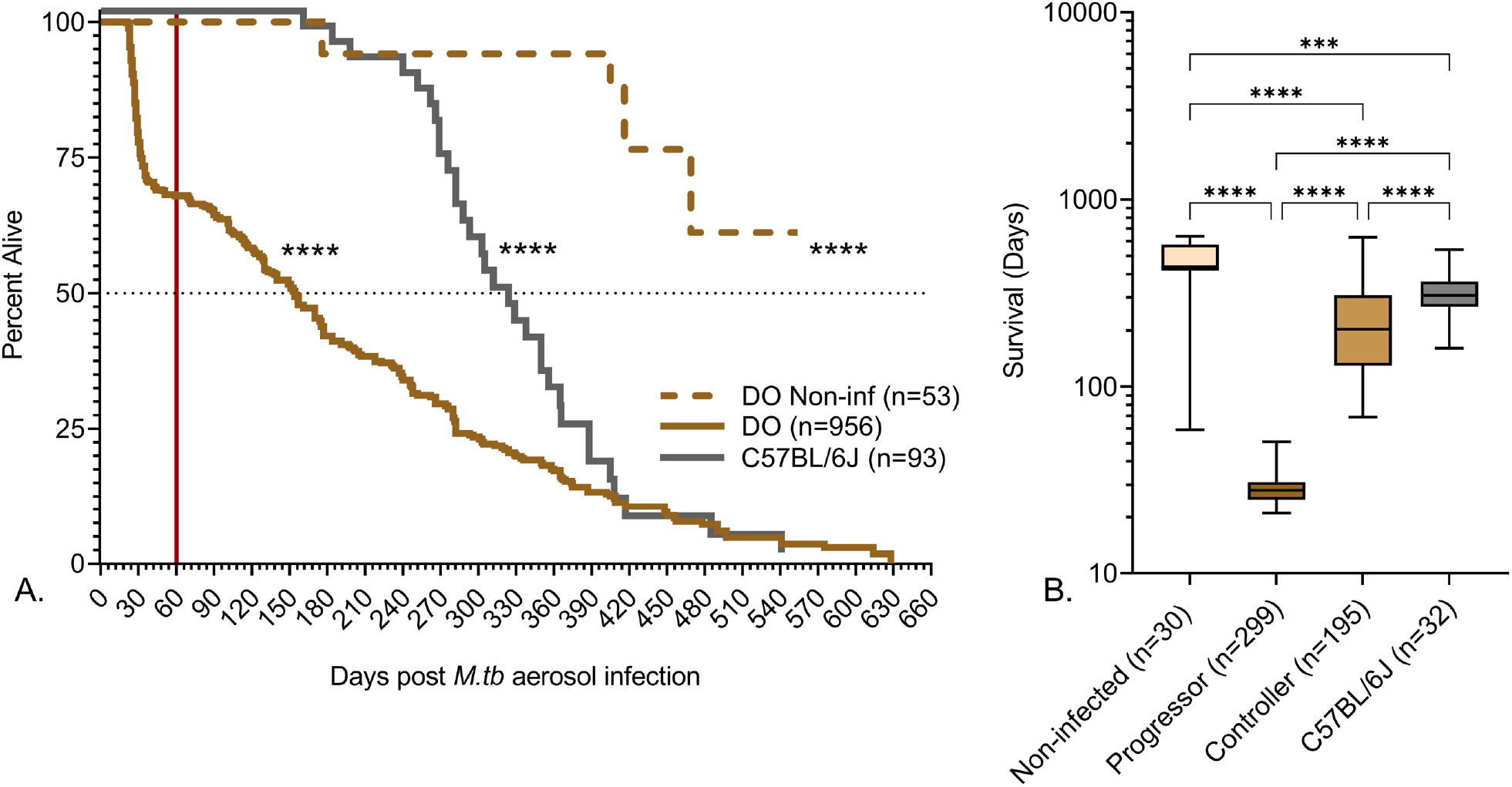
Survival of *M. tuberculosis*-infected mice. We infected 8-10-week-old, female Diversity Outbred (DO) mice and C57BL/6J mice with aerosolized *M. tuberculosis* bacilli and monitored as described in Methods. Non-infected, identically housed, age and gender-matched Diversity Outbred mice served as controls. Mice were euthanized at a predetermined timepoint, or if any one of three morbidity criteria developed: Body condition score <2; severe lethargy; or increased respiratory rate/effort. Panel A shows the percent alive over time (cumulative survival) and the red vertical line marks 60 days of infection when approximately 30% of Diversity Outbred mice succumbed to pulmonary TB and required euthanasia (Progressors). Approximately 70% of Diversity Outbred mice survived longer than 60 days (Controllers). Survival curves for non-infected Diversity Outbred mice (brown dashed line); infected Diversity Outbred (brown solid line); and infected inbred C57BL/6J (solid gray line) mice were significantly different by Mantel Cox log-rank test (**** p<0.0001). Panel B shows significant differences in mean survival days were observed between all classes euthanized due to morbidity attributable to pulmonary TB. A subset of 556 from 1102 mice used in Panel A were available for analysis. Box-and-whiskers plots in panel B show interquartile range with whiskers at the minimum and maximums. Sample sizes are shown in the X-axis and statistical analysis was performed using Brown-Forsythe and Welch’s one-way ANOVA followed by Dunnett’s T3 post-test, **** p<0.0001; ns non-significant.

**Table 1:** Susceptibility to *M. tuberculosis* and forms of TB in humans and Diversity Outbred mice.

|  |  |  |  |  |
| --- | --- | --- | --- | --- |
| **<u>Humans</u>** |  |  |  |  |
| Susceptibility | Highly susceptible | Susceptible | Resistant | Highly resistant |
| Clinical form | Fulminant | Post-primary | Latent Infection | Resistors |
| Histopathologic lesions | Acute, necrotizing | Granulomas, Cavities | Poorly defined | No lesions |
| Survival | Weeks | Years | Decades | Normal lifespan |
| **<u>Diversity Outbred mice</u>** |  |  |  |  |
| Susceptibility | Supersusceptible | Resistant | Resistant | Not observed |
| Clinical form | Progressors | Controllers | Asymptomatic |  |
| Histopathologic lesions | Acute, necrotizing | Chronic, non-necrotizing | Lymphocytic cuffs |  |
| Survival | < 60 days | 60 days to >600 days | At least 60 days |  |

We speculated that resistance to *M. tuberculosis* could have been due to larger body size. However, retrospective analysis of preinfection body weight data failed to identify significant differences (Figure 2A). As expected from inbreeding, age- and gender-matched C57BL/6J mice had a much narrower preinfection weight range, and on average weighed significantly less than Diversity Outbred mice (Figure 2A). Controllers achieved higher body weights (Figure 2B) due to normal growth while infected and longer duration of weight gain (Figure 2C) prior to disease onset. Once disease onset occurred, Controllers lost weight over a longer duration (Figure 2D) and had a slower rate of weight loss or no weight loss (Figure 2F). By euthanasia, both Progressors and Controllers had lost a similar percentage of weight (Figure 2E), and Controllers had a wider range of weight loss. Asymptomatic mice were euthanized before the end of their growth phase and thus achieved lower body weight than non-infected controls (Figure 2B), had a truncated duration of weight gain (Figure 2C), and weight fluctuations without did not lose significant weight (Figures 2D, 2E).

**Figure 2.**
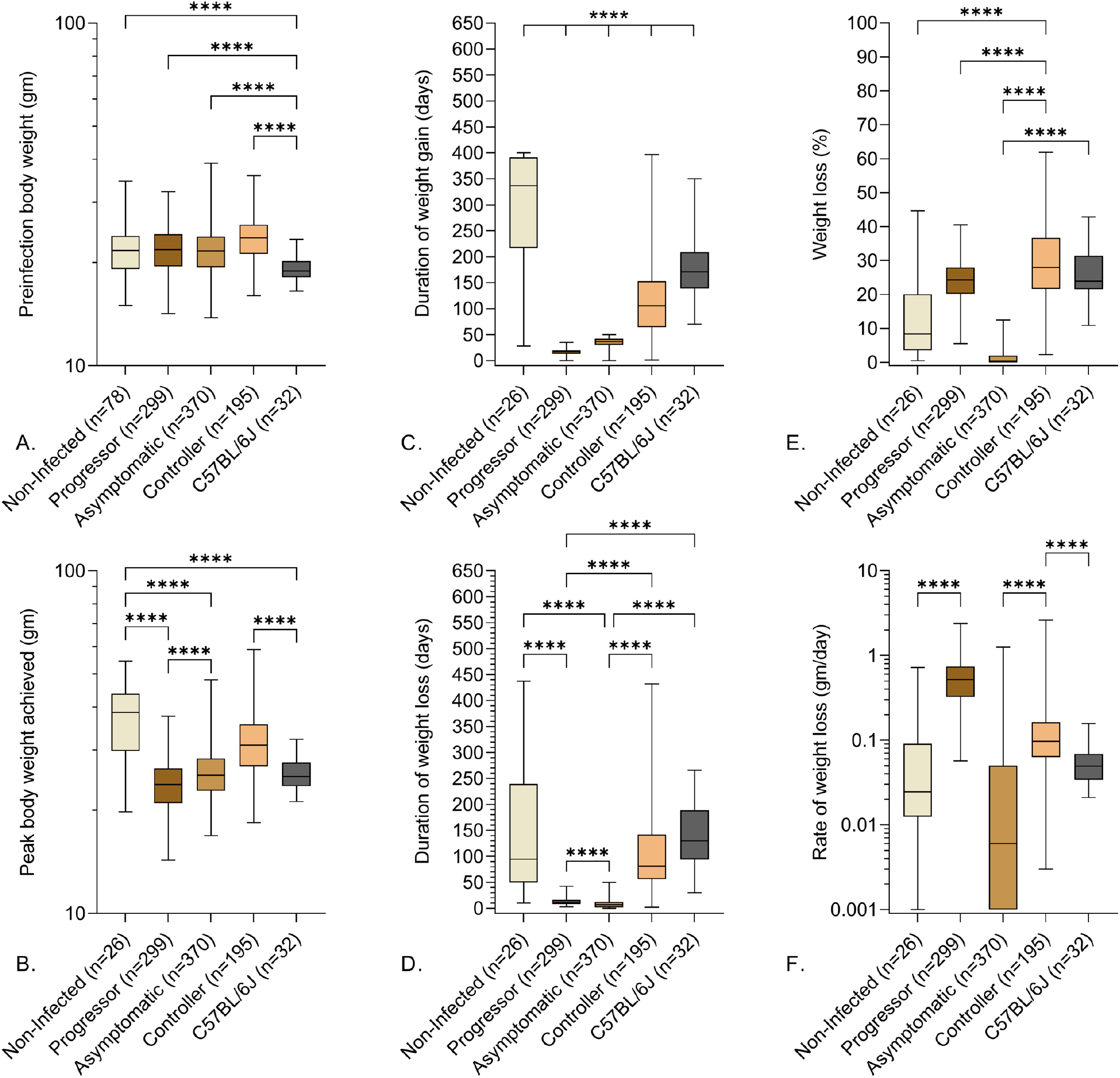
Preinfection body weights and clinical indicators of pulmonary TB in *M. tuberculosis*-infected mice. We infected 8-10-week-old, female Diversity Outbred mice and C57BL/6J mice with aerosolized *M. tuberculosis* bacilli and monitored as described in Methods. Non-infected, identically housed, age and gender-matched Diversity Outbred mice served as controls. Mice were euthanized at a predetermined timepoint, or if any one of three morbidity criteria developed: Body condition score <2; severe lethargy; or increased respiratory rate/effort. (A) Preinfection body weights; (B) Peak body weight achieved during infection; (C) Duration of weight gain; (D) Duration of weight loss); (E) Percent of peak body weight lost; and (F) Rate of weight lost are shown. Box-and-whiskers plots in all panels show interquartile range with whiskers at the minimum and maximum. Sample sizes are shown in the X-axis and statistical analyses were performed using Brown-Forsythe and Welch’s one-way ANOVA followed by Dunnett’s T3 post-test, **** p<0.0001; non-significant p-values are not shown.

We expected Controllers with end-stage chronic pulmonary TB to achieve the same level of *M. tuberculosis* lung burden as Progressors with end-stage acute pulmonary TB. However, this was not true: Controllers had significantly lower *M. tuberculosis* burden than Progressors (Figure 3A). Likewise, we expected Controllers and Progressors to have similar levels of inflammatory mediators. However, this also was not true: the biomarkers we previously identified as diagnostically useful such as matrix metalloproteinase 8 (MMP8), CXCL1, tumor necrosis factor (TNF), and interferon gamma (IFN-γ) [23] were significantly lower in Controllers compared to Progressors (Figures 3B, 3C, 3E, 3F) with the exception of interleukin-10 (IL-10) (Figure 3D). These differences indicate that end-stage pulmonary TB in Diversity Outbred mice has two distinct pathogeneses: an acute form that is highly necrotizing, inflammatory and promotes high *M. tuberculosis* bacillary growth, and a chronic form that is less inflammatory. Asymptomatic mice had significantly lower *M. tuberculosis* burden, CXCL1, and MMP8 and trend for lower levels of TNF, IL-10, and IFN-γ than Progressors. In contrast, only lung *M. tuberculosis* burden and IL-10 were significantly lower between Asymptomatic mice and Controllers (Figures 3A, 3D).

**Figure 3.**
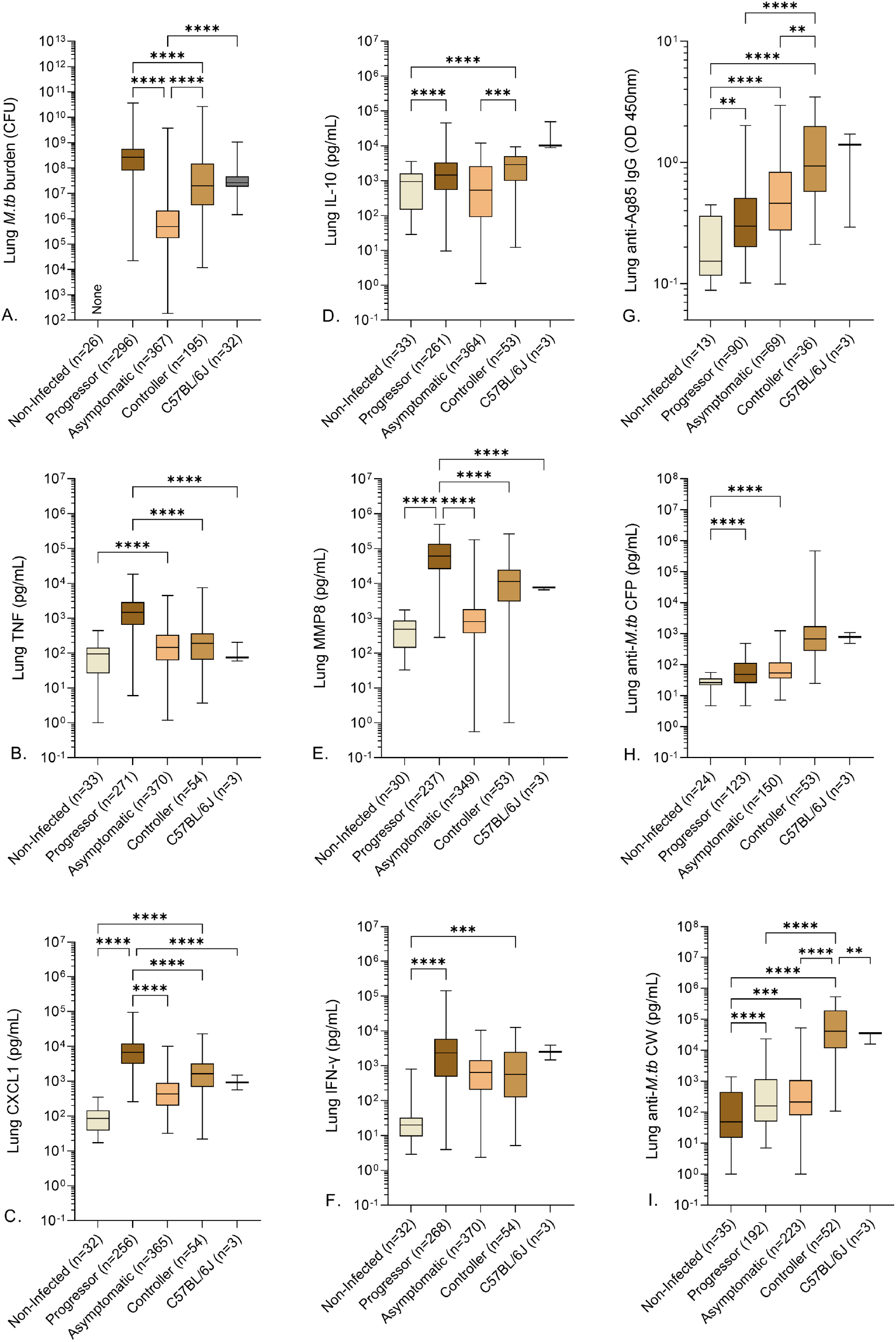
Lung *M. tuberculosis* burden, cytokines, chemokines and anti-*M. tuberculosis* antibodies in *M. tuberculosis*-infected mice. We infected 8-10-week-old, female Diversity Outbred mice and C57BL/6J mice with aerosolized *M. tuberculosis* bacilli and monitored as described in Methods. Non-infected, identically housed, age and gender-matched Diversity Outbred mice served as controls. Mice were euthanized at a predetermined timepoint, or if any one of three morbidity criteria developed: Body condition score <2; severe lethargy; or increased respiratory rate/effort. We quantified *M. tuberculosis* colony forming units in the lungs (A) and measured eight lung proteins using sandwich ELISAs (B-I). Box-and-whiskers plots in all panels show interquartile range with whiskers at the minimum and maximum. Sample sizes are shown in the X-axis. Statistical analyses were performed using Kruskal-Wallis one-way ANOVA with Dunn’s multiple comparisons post-tests (panel A) or Brown-Forsythe and Welch’s one-way ANOVA followed by Dunnett’s T3 post-test (panels B-F) ** p<0.01; ***p<0.001; ****p<0.0001; non-significant p-values are not shown.

As a next step, we analyzed three different modalities: protein measurements, histopathology, and gene-expression profiles with both machine learning and statistical methods (See Figure 4).

**Figure 4:**
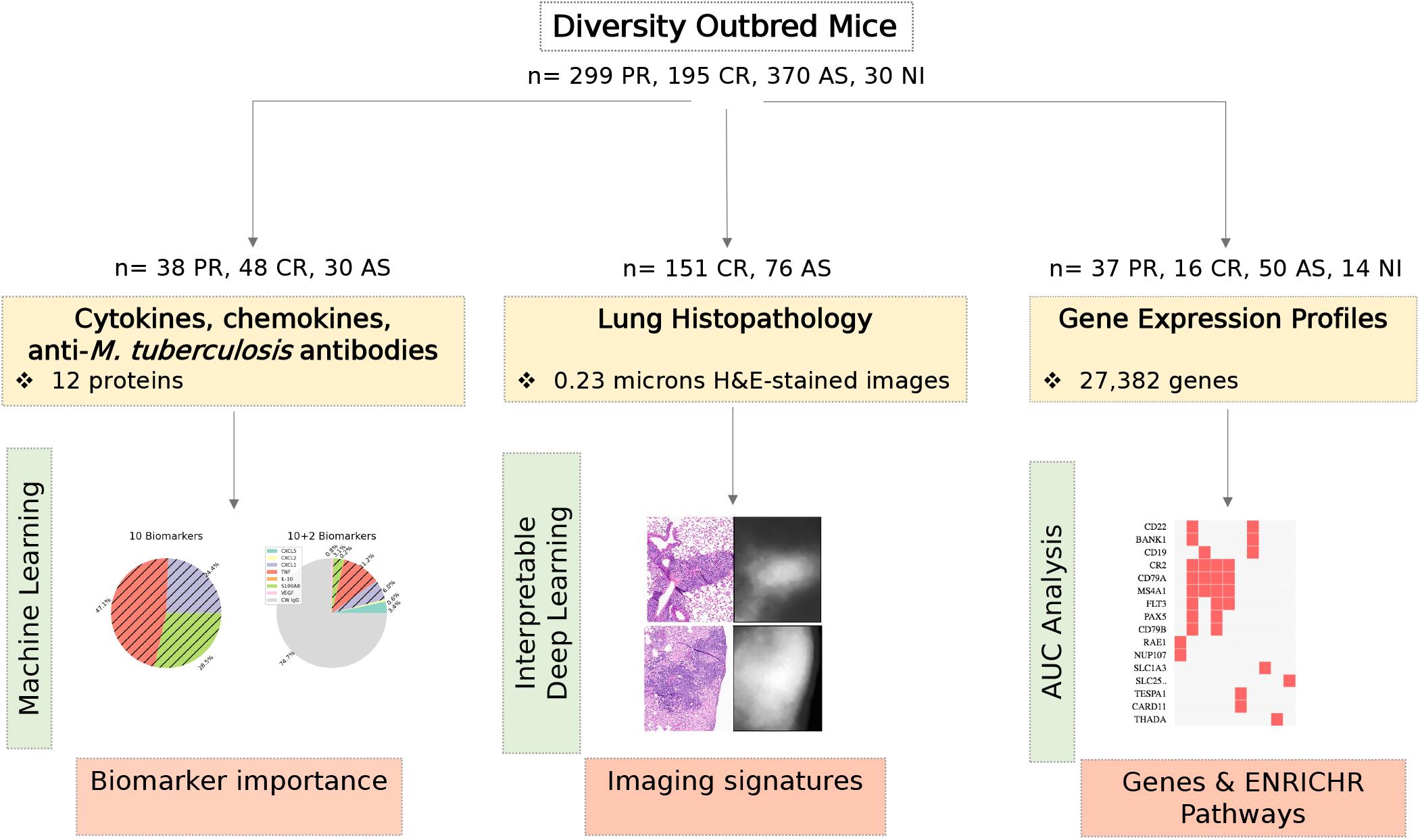
Three modalities: protein measurements, H&E-stained images, and gene expression profiles used for characterizing the resistance to M. tuberculosis in Diversity Outbred mice. We used statistical/machine learning approaches to quantify the feature importance of protein biomarkers, an interpretable deep learning model to identify signature regions of H&E-stained slides, and AUC analysis to filter genes for subsequent ENRICHR pathway analysis.

### A panel of lung cytokines, chemokines, and IgG antibodies classify acute and chronic pulmonary TB but not asymptomatic lung infection

To classify acute vs chronic end-stage pulmonary TB (i.e., Progressors from Controllers) and asymptomatic infection, we analyzed lung proteins (i.e., immune cytokines, chemokines, growth factors), and used them for pairwise comparisons (Table 2). We first trained an L1-regularized logistic regression model using a panel of ten cytokines, chemokines, and growth factors: CXCL5, CXCL2, CXCL1, TNF, IFN-γ, IL-12, IL-10, MMP8, S100A8, and VEGF. The classifier had high 30-fold-cross-validation AUC (0.966) for Progressor vs Asymptomatic (Supplementary Figure 1) but performed relatively poorly (0.792 and 0.803) for comparisons against Controllers (Table 2). When we added measurements of antibodies to the panel of lung proteins (anti-*M. tuberculosis* CW IgG and anti-*M. tuberculosis* CFP IgG), the 30-fold-cross-validation performance improved in all three problems (Table 2). The improvement was the highest for the Progressor vs Controller comparison in which the AUC increased from 0.792 to 0.933 (Figure 5A). That improvement can be attributed to anti-*M. tuberculosis* CW specifically which had the highest average % importance while the other antibody included, anti-*M. tuberculosis* CFP, was not used by the model (Figure 5B). The classification between Controllers and Asymptomatic mice was the most challenging (0.83 AUC) (Table 2, Supplementary Figure 2). When an additional non-linear classifier, Gradient Tree Boosting, was tested the AUC did not improve (Supplementary Table 1). All but one of the six panels performed with >90% accuracy when tested with (n=22) non-infected Diversity Outbred mice previously unused during the training (Supplementary Table 2). One panel, although successful for classifying between the two forms end-stage pulmonary TB using anti-bodies, had low classification accuracy (32%) for the non-infected mice. That is because the non-infected Diversity Outbred mice had low levels of anti-*M. tuberculosis* CW (Figure 3) which the classifier associates with lower survival (Figure 5).

**Figure 5.**
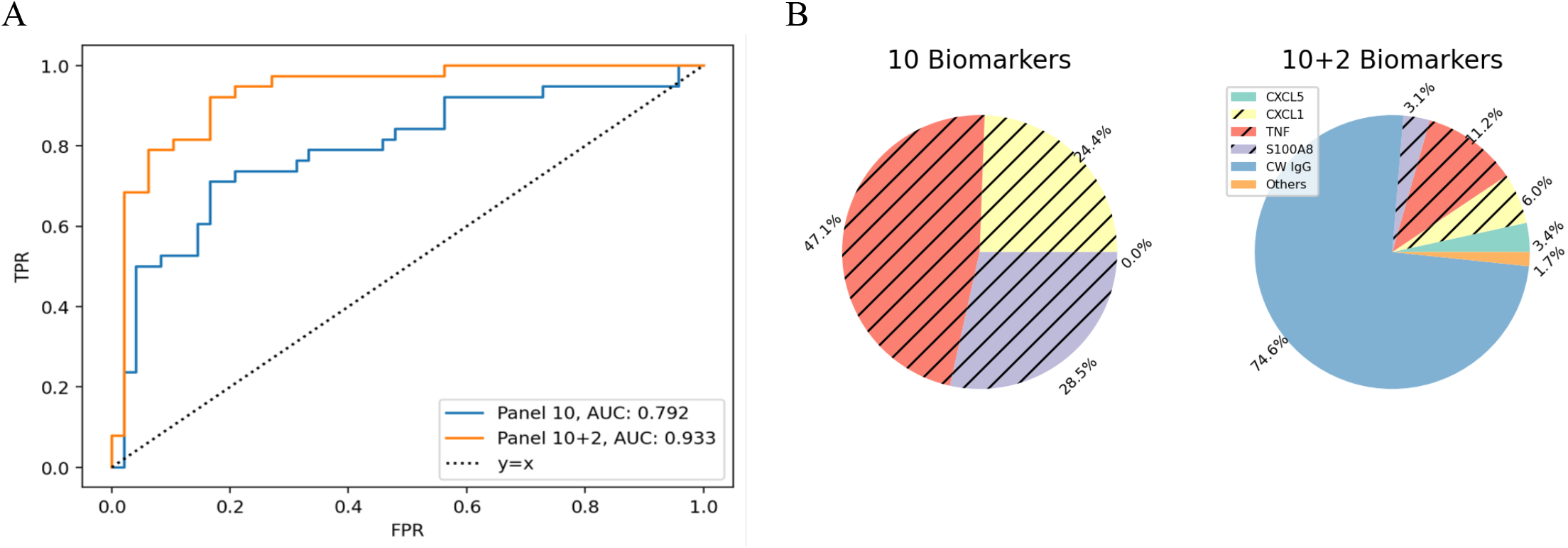
Feature Importance Analysis for classification between Progressors and Controllers using cytokine, chemokine, and anti-*M. tuberculosis* antibodies measurements. a) ROC curve comparison of the ten-biomarker panel (blue) and twelve-biomarker panel (orange) for the Progressor vs Controller comparison. b) Percent Importance of the different biomarkers in two panels. Logistic regression is the classifier and importance scores are averaged over 30 folds. Biomarkers corresponding to the unhatched colors are associated with longer survival and vice versa for the hatched colors. Biomarkers with less than 1% importance are omitted. The feature scores for other comparisons are in Supplementary Figure 1 and Supplementary Figure 2.

**Table 2:**
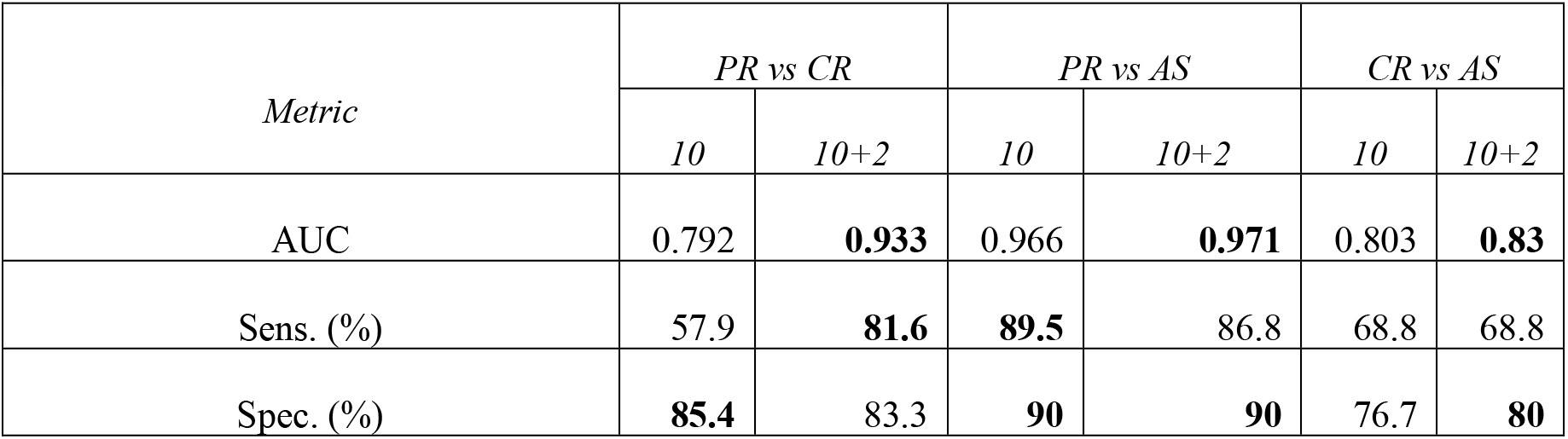
Thirty-fold-cross-validation performance of the two panels (10 & 10+2 antibodies) in three classification tasks (Progressor vs Controller, Progressor vs Asymptomatic, and Controllers vs Asymptomatic). Including antibodies improves the AUC in all three comparisons. For each comparison best result is highlighted in bold. The sensitivity is calculated with respect to Progressors in the first two comparisons and Controllers in the last comparison. Please refer to Supplementary Figure 3 for the confusion matrices. Abbreviations: PR: Progressor, CR: Controller, AS: Asymptomatic.

| *Metric* | *PR vs CR* |  | *PR vs AS* |  | *CR vs AS* |  |
| --- | --- | --- | --- | --- | --- | --- |
|  | *10* | *10+2* | *10* | *10+2* | *10* | *10+2* |
| AUC | 0.792 | **0.933** | 0.966 | **0.971** | 0.803 | **0.83** |
| Sens. (%) | 57.9 | **81.6** | **89.5** | 86.8 | 68.8 | 68.8 |
| Spec. (%) | **85.4** | 83.3 | **90** | **90** | 76.7 | **80** |

### Qualitative evaluation of lung granulomas yields insight

Anti-*M. tuberculosis* cell wall IgG improved the classification of Progressors and Controllers (Figure 5) but not asymptomatic lung infection. To find indicators of asymptomatic lung infection, a board-certified veterinary pathologist (GB) examined lung sections of Progressors, Controllers, and mice with asymptomatic lung infection (Figure 6). Qualitative differences in size, cellular infiltrates and distribution of granulomas were noted, similar to previous publications [18, 23, 24, 38]. The lungs of Progressors contained coalescing necrotizing granulomas with abundant pyknotic nuclear debris within alveoli and bronchioles, and necrosis of alveolar septae (Panel A-D), often with fibrin thrombosis of septal capillaries (not shown). In contrast, the lungs of Asymptomatic mice typically contained small, discrete, non-necrotizing lesions with perivascular and peribronchiolar aggregates of lymphocytes, and few neutrophils (Panels E-H). The lungs of Controllers typically contained diffuse, macrophage-dominated lesions, with many foamy macrophages, dense foci of lymphocytes and plasma cells and occasional multinucleated giant cells (Panels I-L), resembling end-stage pulmonary TB in the commonly used inbred mouse strain C57BL/6J [43]. Additional features in Controllers with end-stage pulmonary TB included cholesterol clefts, small pyogranulomas, septal fibrosis, cavitation with peripheral fibrosis, and bronchiolar obstruction with epithelial degeneration (not shown).

**Figure 6.**
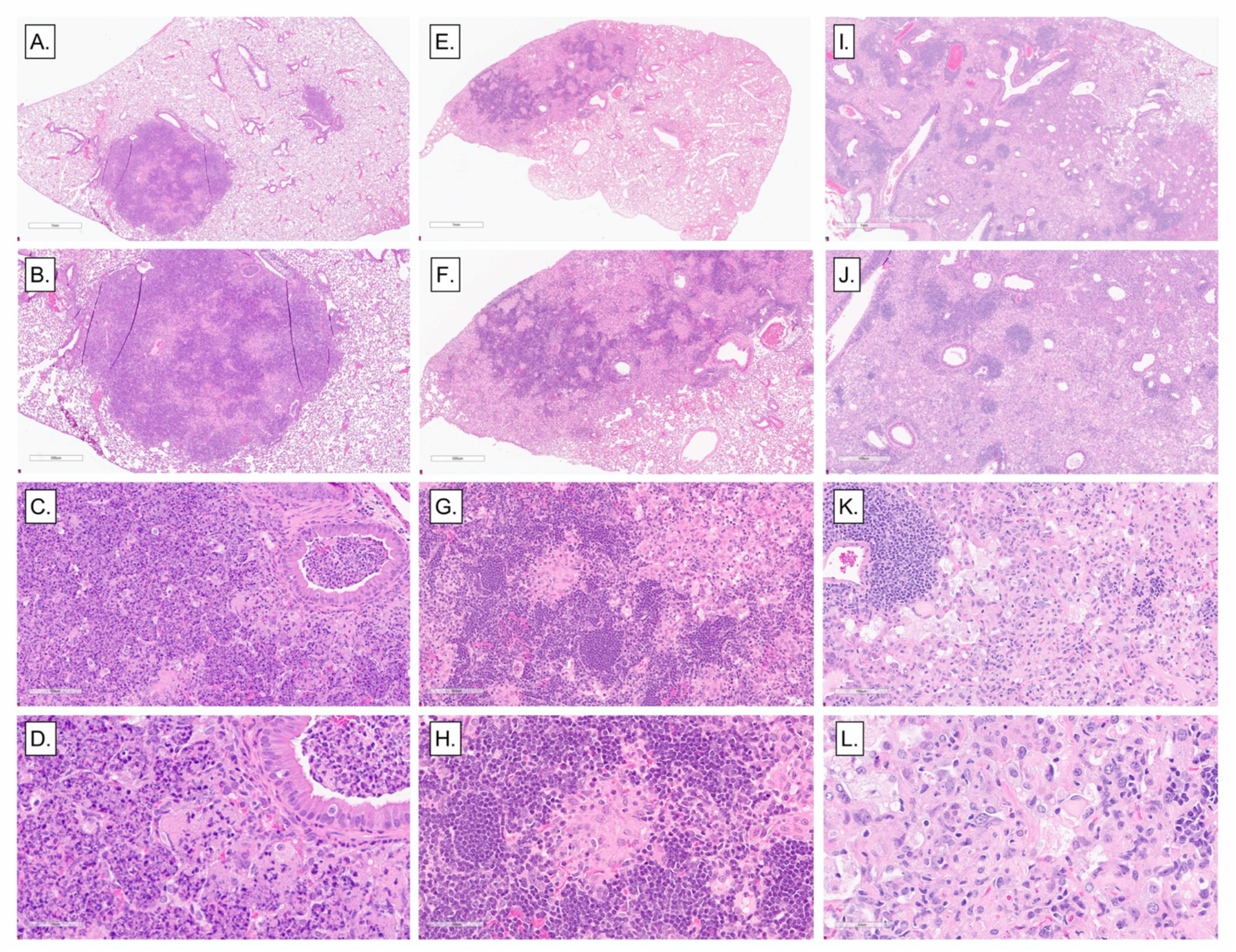
Representative histopathologic lesions in the lungs of *M. tuberculosis-*infected Diversity Outbred mice. We infected 8-10-week-old, female Diversity Outbred mice with *M. tuberculosis* bacilli by inhalation and lung lobes fixed, stained, sectioned for microscopic examination. Representative necrosuppurative lung lesions with bronchiolar obstruction in Progressors are shown in panels A through D; Non-necrotizing lymphohistiocytic lung lesions in Asymptomatic mice are shown in panels E through H. Diffuse, non-necrotizing lesions with abundant macrophages, foamy macrophages and few lymphocytic foci in Controllers are shown in panels I through L. Magnifications are 2x; 4; 20x, and 40x.

### A deep-learning neural network produced an accurate and interpretable imaging biomarker of asymptomatic lung infection

Qualitative histopathological evaluation of granulomas provided some insight; however, quantification of unique histopathological granuloma was not feasible. Therefore, we trained and validated a deep learning neural network using multiple instance learning and attention-based pooling to (i) classify Asymptomatic mice and Controllers; and (ii) identify regions within the granulomas where the model made classification decisions based on feature importance. Table 3 shows the model achieved close to 90% sensitivity and 70% specificity with AUC close to 90%, an improvement over the lung biomarker panel.

**Table 3.**
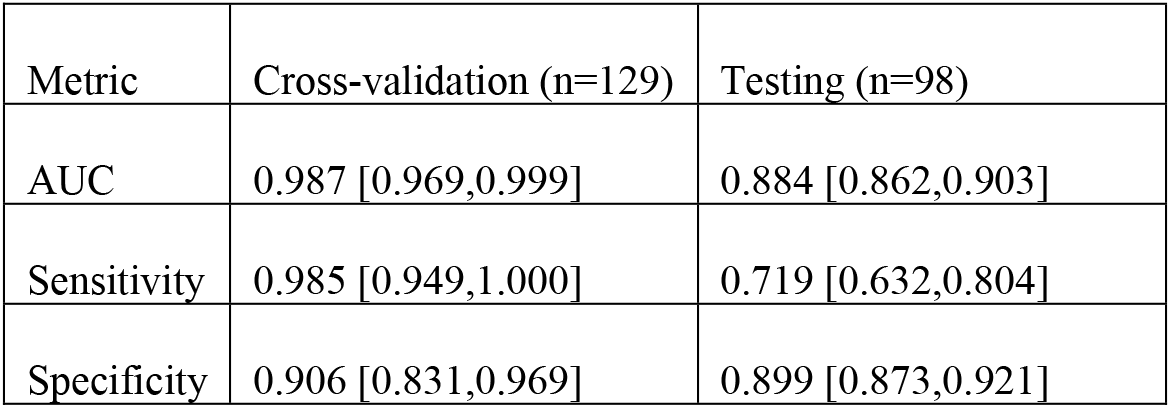
Results of attention-based multiple instance learning model classification of Asymptomatic mice and Controllers.

| Metric | Cross-validation (n=129) | Testing (n=98) |
| --- | --- | --- |
| AUC | 0.987 [0.969,0.999] | 0.884 [0.862,0.903] |
| Sensitivity | 0.985 [0.949,1.000] | 0.719 [0.632,0.804] |
| Specificity | 0.906 [0.831,0.969] | 0.899 [0.873,0.921] |

When we mapped attention weights back to the original images, the granuloma regions used as the basis for classifying asymptomatic lung infection was interpreted by a board-certified veterinary pathologist (GB) as perivascular and peribronchiolar lymphoplasmacytic cuffs (Figure 7a, white areas). In contrast, the neural network did not weight granuloma regions that contained abundant macrophages, cholesterol clefts, or small pyogranulomas (Figure 7b, black areas) which were characteristic of chronic pulmonary TB in Controllers. Overall, the interpretable imaging biomarker results identified perivascular and peribronchiolar lymphoplasmacytic cuffs as key granuloma features of asymptomatic lung infection, which we hypothesized also corresponded to unique functional responses capable of restricting *M. tuberculosis*.

**Figure 7.**
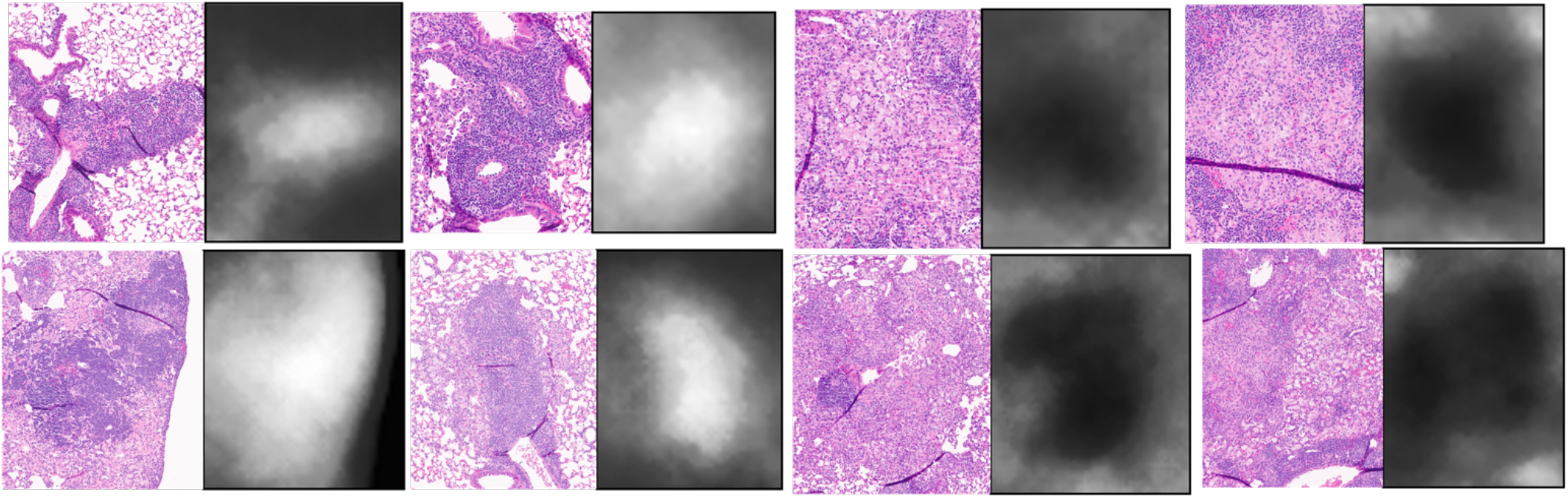
Imaging biomarkers of resistance. The model identified perivascular and peribronchiolar lymphoplasmacytic cuffs as imaging biomarkers for Asymptomatic mice (a), whereas abundant macrophages, cholesterol clefts, or small pyogranulomas were more characteristic of Controllers and represent pulmonary TB disease progression (b).

### Gene expression analysis

To identify functional correlates of perivascular and peribronchiolar lymphoplasmacytic cuffs in asymptomatic lungs, we used transcriptional profiling and pathway analyses on two available data sets. One dataset consistent of n=5 non-infected; n=13 Asymptomatic; n=16 Controller, and n=10 Progressor Diversity Outbred mice. The second dataset consistent of n=9 Non-Infected; n=37 Asymptomatic; and n=27 Progressor Diversity Outbred mice.

Within each dataset, we performed a one-sided AUC analysis (see Methods) comparing the Progressor and Asymptomatic lungs, which identified sets of 2,569 and 6,891 genes expressed at significantly (FDR *q* < 0.05) higher levels in the lungs of Asymptomatic mice within dataset 1 and 2, respectively. These two sets contained 2,264 genes in common, with the average AUC values of the two datasets ranging from 0.743 to 0.969 with the median 0.844 (Supplementary File 1). This set of genes was input using Enrichr for pathway analysis, which identified 33 statistically significant (adjusted *p* < 0.05) Gene Ontology terms (Supplementary Table 3).

In the same manner, we compared Controller and Progressor lungs using dataset 1, and identified a set of 303 genes expressed at significantly (FDR *q* < 0.05) higher levels in the lungs of Controllers. The AUC of the selected genes ranged from 0.888 to 1 with the median 0.919 (Supplementary File 1). Enrichr analysis identified five pathways containing statistically significant, highly expressed genes in Controllers and the pathways represent adaptive immunity. i.e., T cell activation and regulation, interleukin (IL)-21 and IL-21 receptor signaling, and antigen receptor-mediated signaling pathways (Table 4).

**Table 4.** Enrichr analysis resulting from the significantly different highly expressed genes in the lungs of Controllers with end-stage chronic pulmonary TB vs Progressors with end-stage acute pulmonary TB. Only the significant pathways (adjusted *p*<0.05) are displayed.

| Term | Adjusted<br>P-value | Genes |
| --- | --- | --- |
| regulation of lymphocyte activation (GO:0051249) | 0.002 | KAT2A;LCK;SIT1;IKZF3;LAT |
| regulation of T cell activation (GO:0050863) | 0.004 | CD2;KAT2A;SIT1;LCK;TNFSF8;LAX1;LAT |
| cellular response to interleukin-21 (GO:0098757) | 0.044 | IL21R;STAT4;IL2RG |
| interleukin-21-mediated signaling pathway (GO:0038114) | 0.044 | IL21R;STAT4;IL2RG |
| T-helper cell lineage commitment (GO:0002295) | 0.044 | SPN;IRF4;LY9 |

None of the genes were significant at FDR *q* < 0.05 for Asymptomatic vs Controller lungs. Upon further inspection, however, we observed that genes with high diagnostic potential for the binary classification between Asymptomatic and Controller groups can include genes that are elevated in the non-infected lungs (Supplementary Figure 6).

To focus on finding unique functional signatures, we compared asymptomatic lungs with those from all other groups: Controller, Progressor, and non-infected using dataset 1. This identified 105 genes that were expressed at significantly (FDR *q* < 0.05) higher levels in lungs (Figure 8). The AUC values of the identified genes ranged from 0.844 to 0.963 with the median 0.864 (Supplementary File 1). Pathway analysis using Enrichr identified eleven statistically significant (adjusted *p* < 0.05) GO terms associated with the 105 genes (Table 5). Six pathways are specific to B-cell functions. Two pathways indicate generic lymphocyte proliferation and activation, which include B-cell genes (e.g., CD79A). Two indicate positive regulation of antigen-receptor mediated signaling; which include B-cell specific markers and transcription factors for immunoglobulin genes (e.g., CD19, ELF1, PAX5). Overall, 10 of the 11 pathways with highly expressed genes indicate that B cell differentiation, proliferation, activation, and effector functions may be important for establishing asymptomatic lung infection and early resistance to *M. tuberculosis.* The last pathway containing highly expressed genes involved functions of RNA polymerase. Taken together, these results suggest that B cell differentiation, proliferation, activation, and effector functions are important to restrict *M. tuberculosis* growth and establish asymptomatic lung infection.

**Figure 8:**
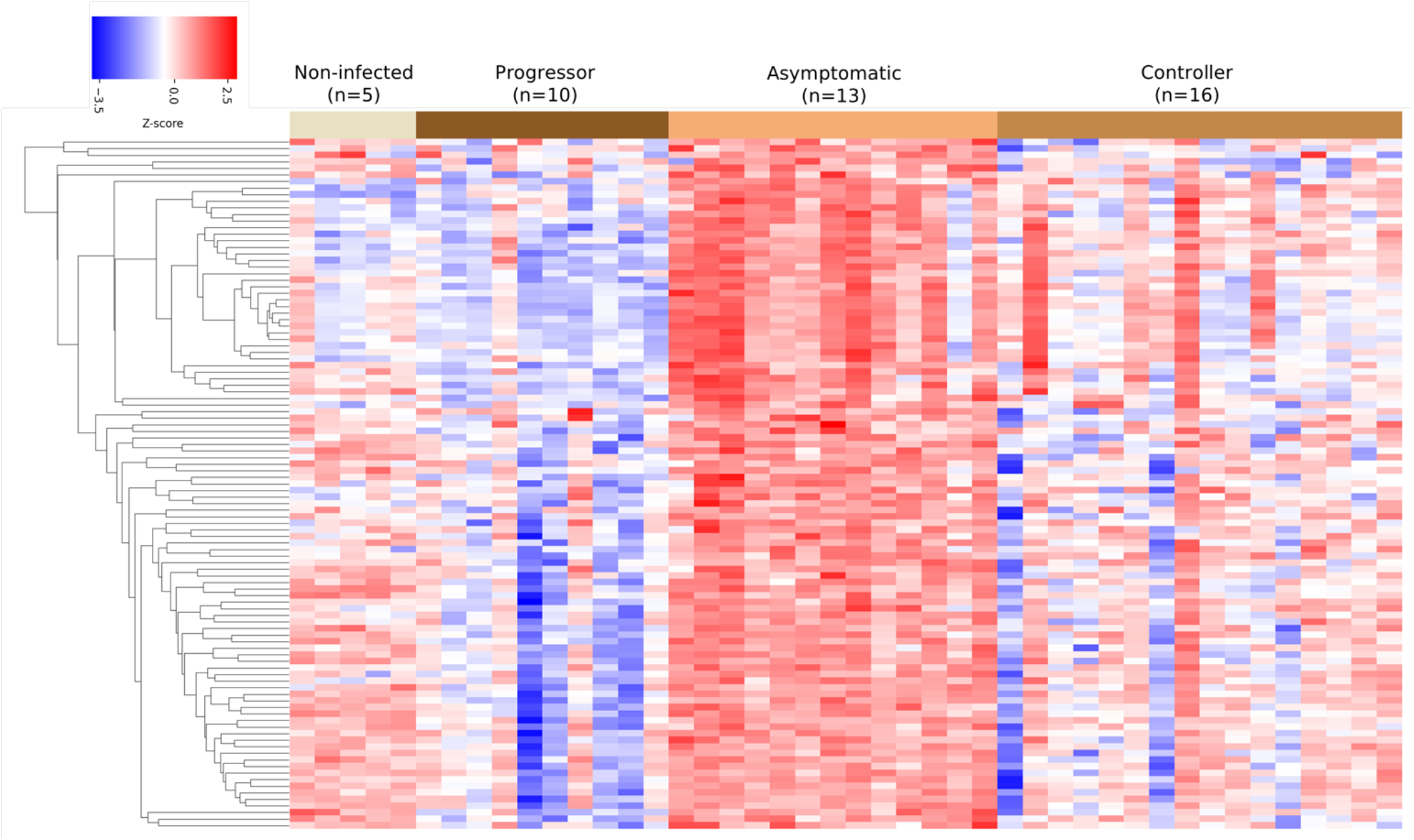
Cluster heat map of the 105 genes selected for the Asymptomatic vs rest classification. Columns correspond to the (n=44) mice used in the Asymptomatic vs rest classification. Rows correspond to the identified 105 genes. Color indicates the z-score and is calculated by first subtracting the row wise average from the gene expression value and next dividing it to the row wise standard deviation. Gene expression values are first log2 transformed.

**Table 5.** Enrichr analysis resulting from the genes selected for the Asymptomatic lung classification vs other groups (Progressor, Controller, non-infected). Only the significant pathways (adjusted *p*<0.05) are shown.

| Term | Adjusted<br>P-value | Genes |
| --- | --- | --- |
| B cell activation (GO:0042113) | <0.000 | FCRL1;CD79B;CD79A;CR2;FLT3;BANK1;MS4A1;CD22 |
| regulation of B cell receptor signaling pathway (GO:0050855) | <0.000 | ELF1;CD19;STAP1;PAX5;CD22 |
| B cell differentiation (GO:0030183) | 0.004 | CD79B;CD79A;CR2;FLT3;MS4A1 |
| B cell proliferation (GO:0042100) | 0.004 | CD79A;CR2;CD19;MS4A1 |
| B cell receptor signaling pathway (GO:0050853) | 0.004 | CD79B;CD79A;CD19;MS4A1 |
| regulation of antigen receptor-mediated signaling pathway (GO:0050854) | 0.004 | ELF1;CD19;PAX5 |
| lymphocyte proliferation (GO:0046651) | 0.004 | CD79A;CR2;FLT3;MS4A1 |
| lymphocyte differentiation (GO:0030098) | 0.007 | CD79B;CD79A;CR2;FLT3;MS4A1 |
| positive regulation of antigen receptor-mediated signaling pathway (GO:0050857) | 0.013 | TESPA1;STAP1;CARD11 |
| transcription-dependent tethering of RNA polymerase II gene DNA at nuclear periphery (GO:0000972) | 0.026 | NUP107;RAE1 |
| regulation of B cell activation (GO:0050864) | 0.026 | BANK1;CD19;CD22 |

## DISCUSSION

Although pulmonary TB remains a global health problem with high morbidity and mortality, ∼90% of humans are highly resistant to infection and have no clinical signs of disease [1, 6, 44-47], making the mechanisms of resistance difficult to identify. A growing body of evidence shows that Diversity Outbred mouse responses to *M. tuberculosis* model humans with similarities in biomarkers of pulmonary TB, and gene expression signatures, providing translational diagnostics and insight into disease pathogenesis [14-18, 20, 23, 30, 48]. With few exceptions, the studies did not define key features of lung resistance to *M. tuberculosis*, which occur in hosts with asymptomatic infection. The novelty of our work here is the approach to find unique features of lung resistance to *M. tuberculosis,* by using the Diversity Outbred mouse population.

To discover signatures of resistance to *M. tuberculosis* infection, we performed long term survival studies in the Diversity Outbred mouse population, and showed survival was bimodal: less than 60 days or greater than 60 days. Interestingly, no infected mice reached the median survival non-infected mice and even the most resistant Diversity Outbred mouse eventually succumbed to chronic progressive pulmonary TB, as in inbred strains [43]. A panel of ten biomarkers which accurately discriminated Progressors with acute pulmonary TB from asymptomatically infected non-progressors [23] could not distinguish Progressors from Controllers that succumbed to chronic pulmonary TB months to > 1 year later. Accurate classification between acute and chronic pulmonary TB was possible when the panel included anti-*M. tuberculosis* cell wall IgG. None of cytokines, chemokines, growth factors or antibodies resulted in a biomarker panel or key feature that could accurately classify Diversity Outbred mice with asymptomatic lung infection. This finding is consistent with the historical challenges to accurately diagnose humans with asymptomatic latent TB infection by using skin tests, interferon gamma responses, or antibody-based serological tests [49-55]. Although, recent studies suggest certain antibodies may provide more diagnostic value [56-58].

Histopathological analyses combined with gene expression was much more useful to find key features of asymptomatic lung infection that have diagnostic value and provide mechanistic insight. Our deep learning model automatically identified a granuloma feature specific to asymptomatic *M. tuberculosis* lung infection, which was interpretable by a pathologist as perivascular and peribronchiolar lymphocytic cuffs. This histological feature aligns with a large body of prior work in inbred mice, non-human primates, and natural experiments in humans (e.g., humans with acquired immune deficiency, or with genetic immune deficiencies) indicating that CD4 T cells and their effector molecules are required for resistance to *M. tuberculosis* [59]. We therefore expected that lung tissue from asymptomatically infected Diversity Outbred mice would contain highly expressed genes and gene expression pathways indicative of CD4 T cell functions. However, we were surprised to find that the differentially expressed genes in lungs of Diversity Outbred mice with asymptomatic *M. tuberculosis* infection corresponded to B-cell functions and signaling, not to CD4 T cell functions and signaling. Of the eleven upregulated pathways in lungs of asymptomatically infected Diversity Outbred mice, ten of the pathways involved B-cell differentiation, proliferation, activation, and effector functions. None of the significant pathways were specific for T cells or CD4 T cells of any subtype.

We compared our gene expression results with a 2020 study by Ahmed *et. al.* [20] which used RNAseq to identify differences between Diversity Outbred mice with a high-risk disease score (n=16), low-risk disease score (n=13), and non-infected (n=10). Of the 105 genes we identified in the lungs of asymptomatically infected Diversity Outbred mice, 14 were reported by Ahmed *et. al*. as significantly overexpressed low-risk vs high-risk disease scores and 52 were significantly overexpressed in the low-risk disease score vs non-infected [20]. The mismatch between our and their findings may reflect differences in methods (microarray vs RNAseq) and sample size (hundreds vs tens). More importantly, among the matching sets of 14 and 52 highly expressed genes, 10 overlapped: *Pax5, Zfp318, Thada, Ralgps2, Dclk2, Itpr2, Cyb561a3, Dock8, F8*, and *Bach2.* Many of these genes transcriptionally regulate B-cell differentiation, and immunoglobulin production. Esaulova *et. al.* reported related findings in a 2021 study that B cell follicles were smaller in the lungs of macaques with pulmonary TB (n=5) compared to those with latent TB infection (n=2) and the negative correlation between the B cell follicle size and lung *M. tuberculosis* burden supported a protective role of inducible Bronchiolar Associated Lymphoid Tissue [60]. Although scRNA-seq analysis indicated the relative number of CD79A+ B cells was higher in the pulmonary TB compared latent TB infection [60], our results suggest the opposite: that lungs of asymptomatically infected Diversity mice had significantly higher expression of CD79A.

Our studies examining resistance to *M. tuberculosis* using protein biomarkers, lung histopathology, deep-learning neural networks, and gene expression profiles provide substantial insight into the mechanisms of disease and resistance, there are limitations. Although we have a nearly 900 mice available for survival and weight analysis, we have gaps in the data set, and subsequent analyses used data sets including slightly over 100-200 samples. In part these data gaps reflect challenges with the small size of mouse lungs, and the samples have limited volumes for example in biomarker panel studies. The gene expression profiles had 117 samples available, and second sets of gene expression profiles remain in processing and will become available for future studies. Secondly, this work did not explore joint analysis of different modalities because using either gene expression profiles or histopathology slides alone enabled accurate classification between acute pulmonary TB in Progressors, chronic pulmonary TB in Controllers and the lungs from asymptomatically infected mice. A future direction can be capturing the intermodal relationships for identifying more complex resistance signatures and boosting the diagnostic accuracy.

Overall, our results show two main findings that have important implications. First, there are two distinct forms of end-stage pulmonary TB in Diversity Outbred mice, which can inform the pathogenesis of pulmonary TB in humans, and additionally support research to discover and host-directed therapies against these two forms of TB. Second, by applying novel computational approaches, image analysis and lung transcriptional profiles, we found granuloma regions of perivascular and peribronchiolar cuffs specific to asymptomatic lung infection and lung functional responses, which show B-cells may be more important in establishing asymptomatic *M. tuberculosis* lung infection. Future studies using the Diversity Outbred mouse population will define the genetic control upstream of these B-cell responses to *M. tuberculosis* infection to understand how host genotype influences outcomes to *M. tuberculosis* infection.

## SUPPLEMENTARY FIGURES AND TABLES

**Supplementary Figure 1.**
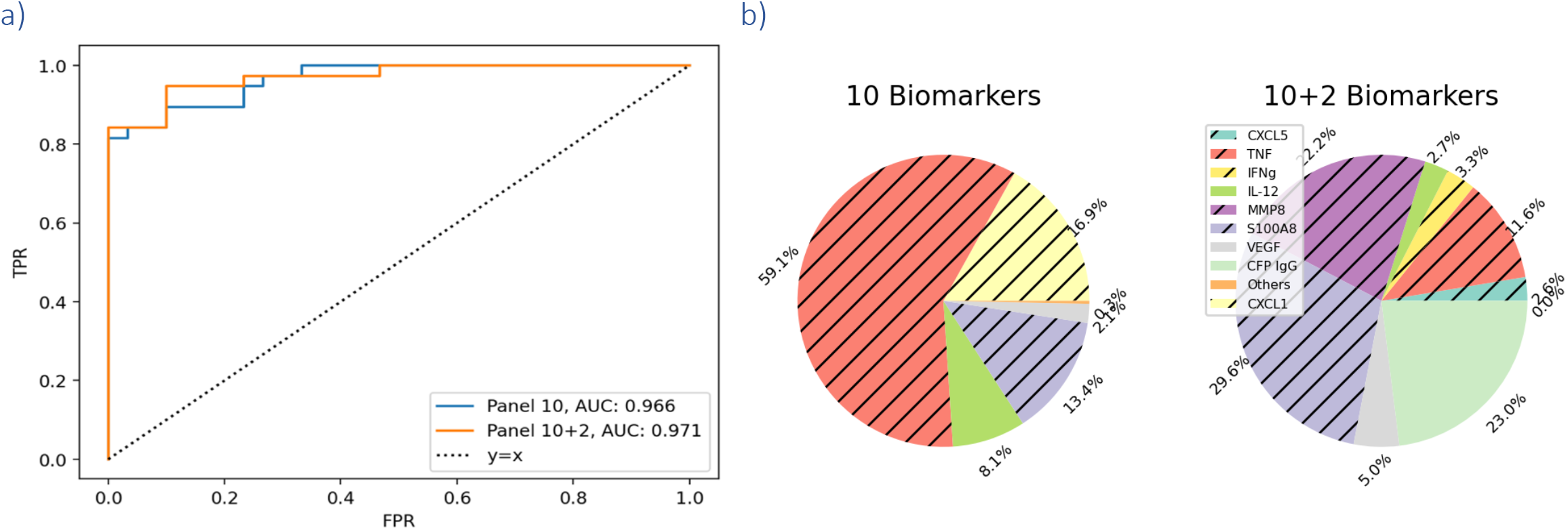
a) ROC curve comparison of the ten-biomarker panel (blue) and twelve-biomarker panel (orange) for the classification between Progressors and Asymptomatic mice. b) Percent Importance of different biomarkers for classifying between Progressors and Asymptomatic mice. Logistic regression is the classifier and importance scores are averaged over 30 folds. Biomarkers corresponding to the unhatched colors are associated with longer survival and vice versa for the hatched colors. Biomarkers with less than 1% importance are omitted.

**Supplementary Figure 2:**
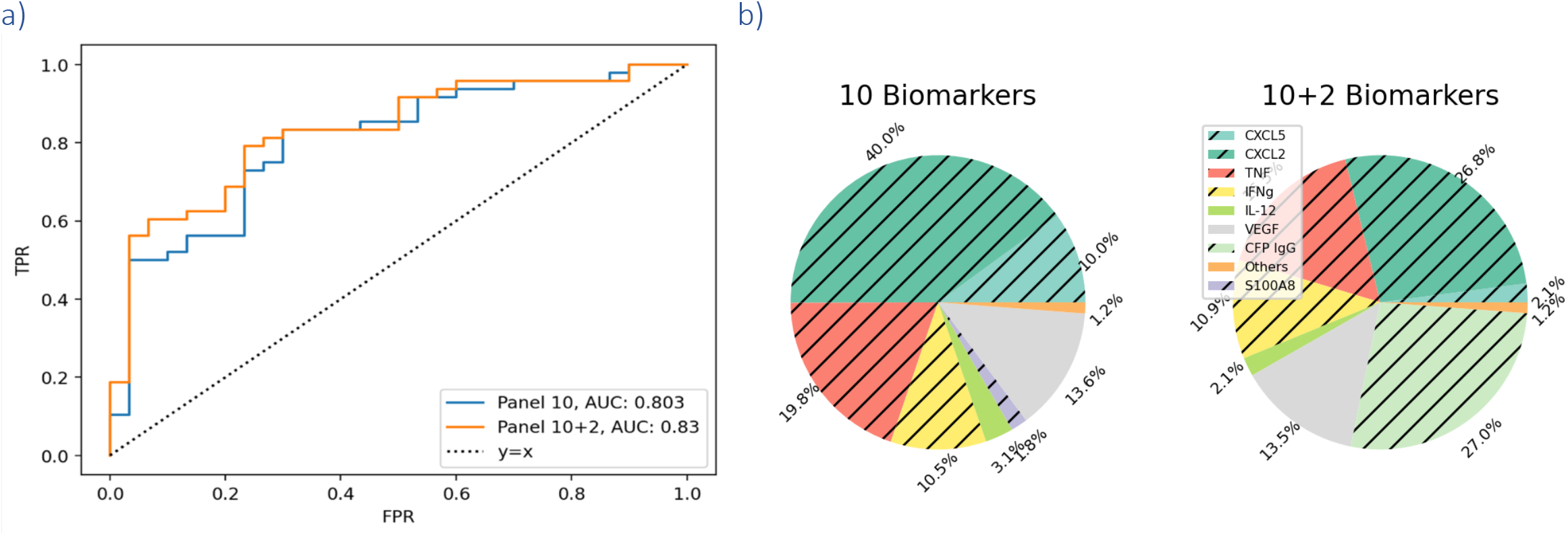
a) ROC curve comparison of the ten-biomarker panel (blue) and twelve-biomarker panel (orange) for the classification between Controllers and Asymptomatic mice. b) Percent Importance of different biomarkers for classifying between Controllers and Asymptomatic mice. Logistic regression is the classifier and importance scores are averaged over 30 folds. Biomarkers corresponding to the unhatched colors are associated with longer survival and vice versa for the hatched colors. Biomarkers with less than 1% importance are omitted.

**Supplementary Figure 3:**
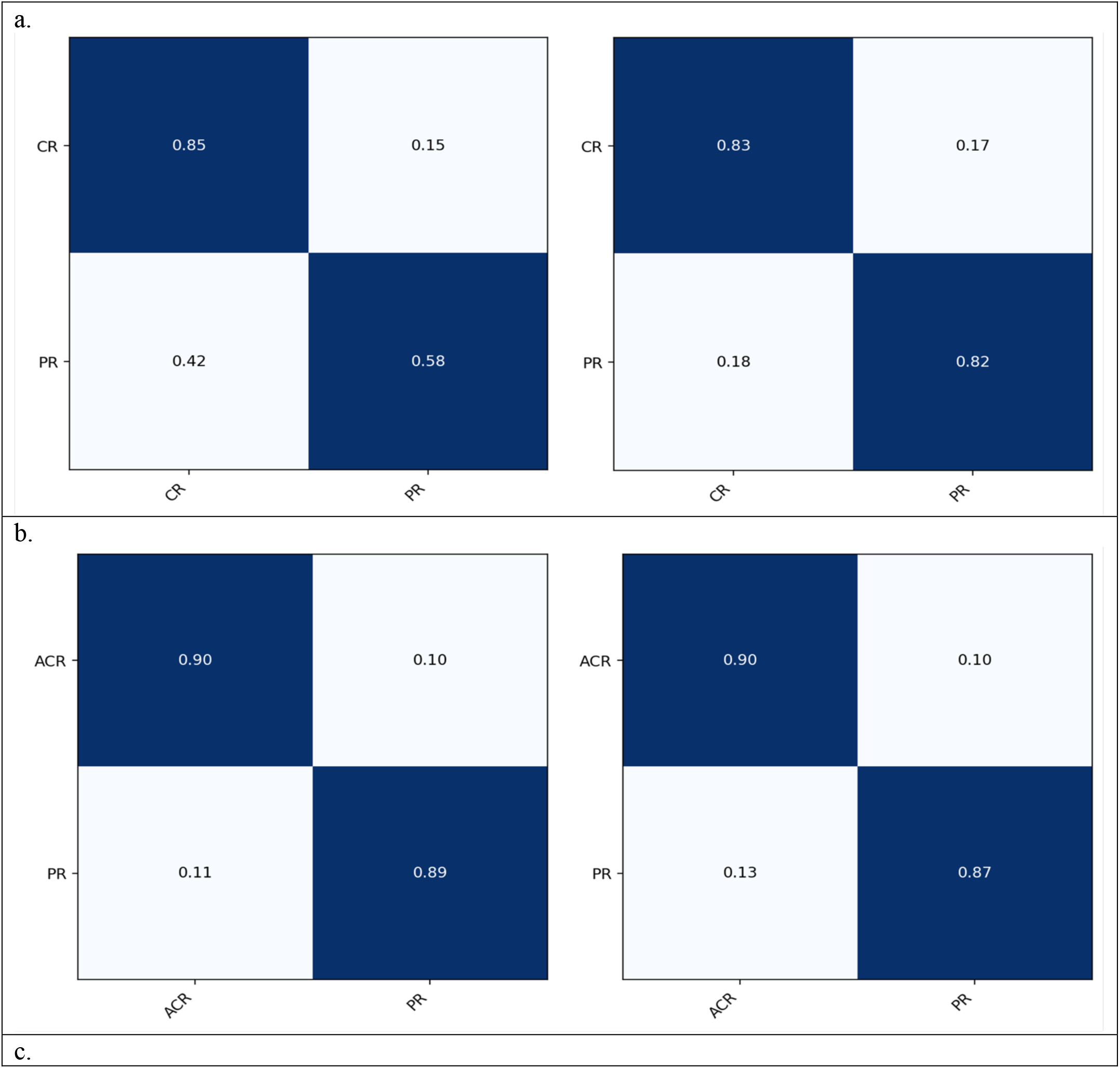
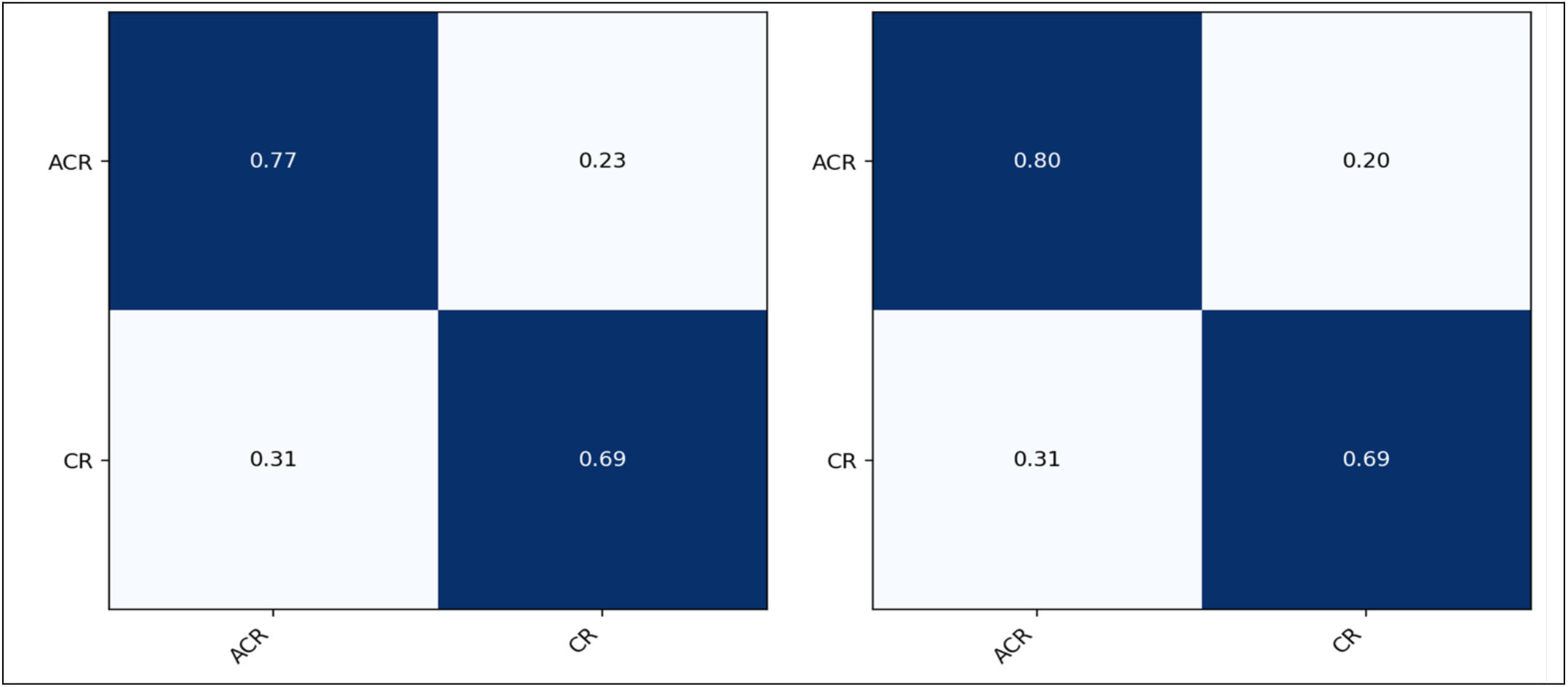
Confusion matrices for three different classification tasks: a. Controllers vs Progressors, b. Asymptomatic mice vs Progressors and c. Asymptomatic mice vs Controllers. For each classification task, we display two confusion matrices corresponding to results of panels with 10 (left) and 12 (right) biomarkers respectively. PR: Progressors, CR: Controllers, ACR: Asymptomatic mice.

**Supplementary Table 1:** Performance of the two panels (10 & 10+2 antibodies) in Controllers (CR) vs Asymptomatic mice (AS) using Xgboost. For each comparison best result is highlighted in bold. The positive class is CR. Confusion matrices are given.

|  |  |  |
| --- | --- | --- |
| *Metric* | *CR vs AS* |  |
|  | *10* | *10+2* |
| AUC | 0.77 | **0.774** |
| Sens. (%) | 72.9 | 72.9 |
| Spec. (%) | **80.0** | 76.7 |

**Supplementary Figure 4:**
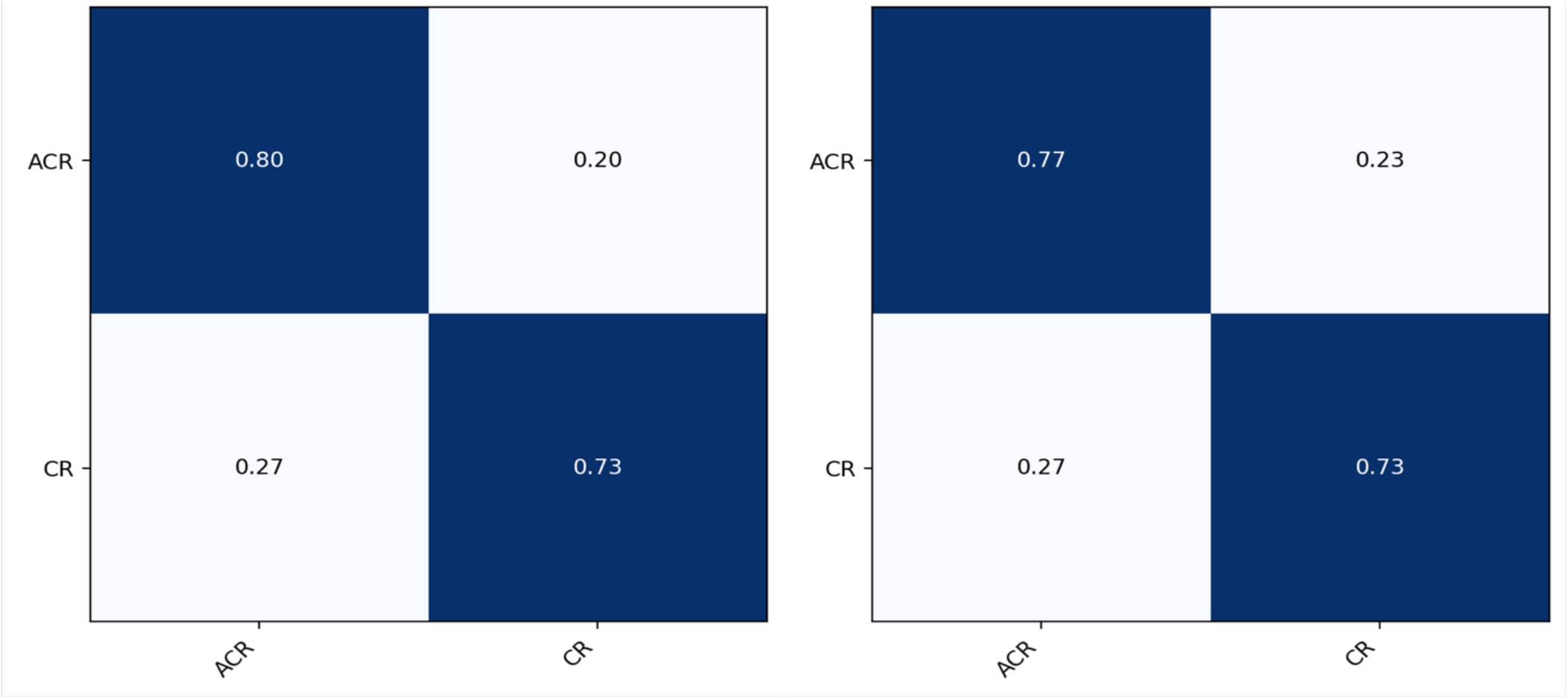
Confusion matrices corresponding to the Xgboost model for Asymptomatic vs Controller comparison. Left and right columns correspond to panels with 10 and 12 biomarkers respectively. CR: Controllers, ACR: Asymptomatic mice.

**Supplementary Table 2:** Non-infected mice classification performance of the two panels (10 & 10+2 antibodies) trained on three classification problems (Progressors vs Controllers, Progressors vs Asymptomatic mice, and Controllers vs Asymptomatic mice). All n=22 non-infected mice with complete data are used and none of the non-infected mice are used during model training. A prediction is deemed correctly classified if it is predicted as the negative class. Last row denotes the confidence of the model i.e. the average probability of a non-infected mice belonging to the negative class. It indicates the non-infected samples are close to the decision boundary of the panel with the least classification accuracy (31.8%). PR: Progressors, CR: Controllers, AS: Asymptomatic mice, NI: Non-infected mice.

| *Metric* | *Trained for*<br>*PR vs CR* |  | *Trained for*<br>*PR vs AS* |  | *Trained for*<br>*CR vs AS* |  |
| --- | --- | --- | --- | --- | --- | --- |
|  | *10* | *10+2* | *10* | *10+2* | *10* | *10+2* |
| Accuracy for NI mice (%) | 100 | 31.8 | 95.5 | 90.9 | 90.9 | 90.9 |
| Avg. Prob. (%) | 68 | 45.1 | 84.2 | 70.3 | 62.6 | 65.8 |

**Supplementary Figure 5.**
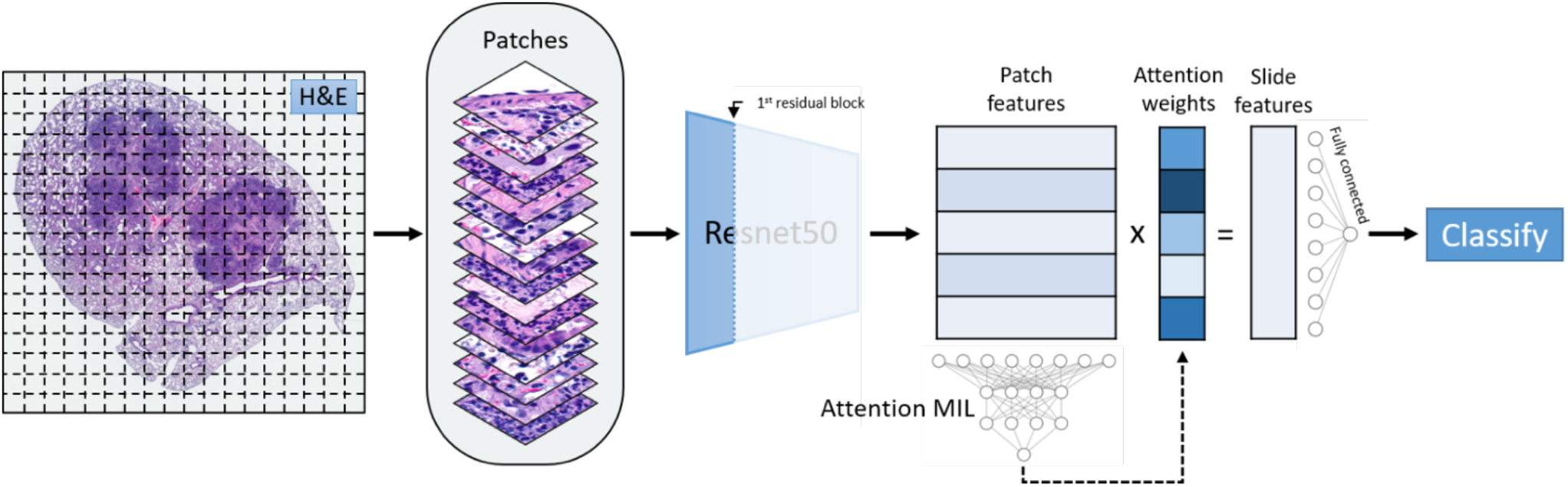
Model overview. Each embedded instance is passed through a shallow feature extractor consisting of two fully connected layers with ReLU activation. Then, each instance is passed through the attention mechanism. This consists of two parallel fully connected layers, a dot product between their outputs, then a final fully connected layer to yield an attention weight. Attention weights are scaled using softmax then dotted with respective outputs from the shallow feature extractor. The resulting slide-level feature vector is then classified into either controller or asymptomatic category.

**Supplementary Figure 6:**
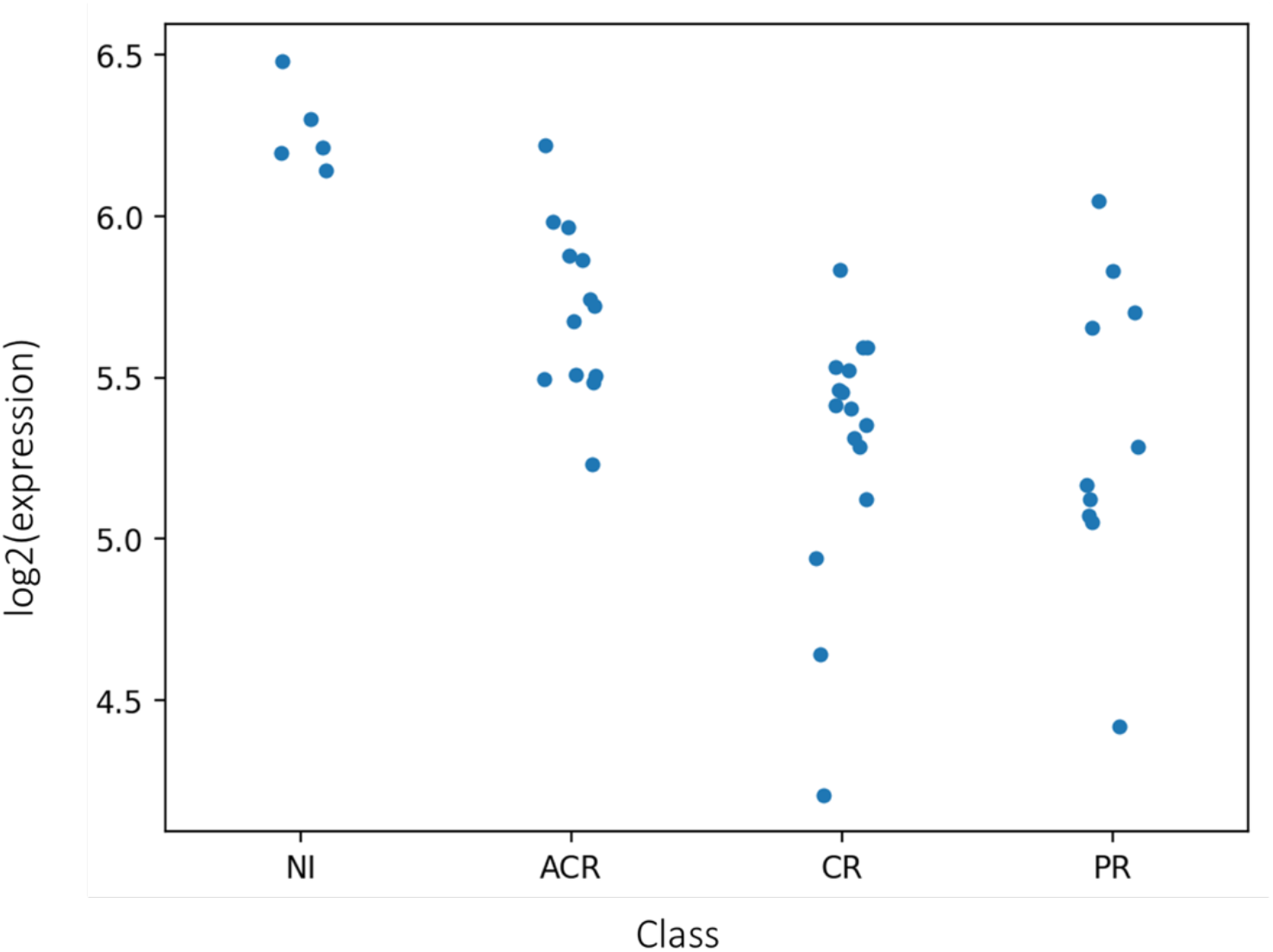
Log2 expression of gene F2rl1. Although F2rl1 had high AUC (0.83) for classifying between Asymptomatic and Controller, expression of the gene was lower in asymptomatic mice compared to the non-infected Diversity Outbred mice. NI: non-infected mice, ACR: asymptomatic mice, CR: controllers, PR: progressors.

**Supplementary Table 3:**
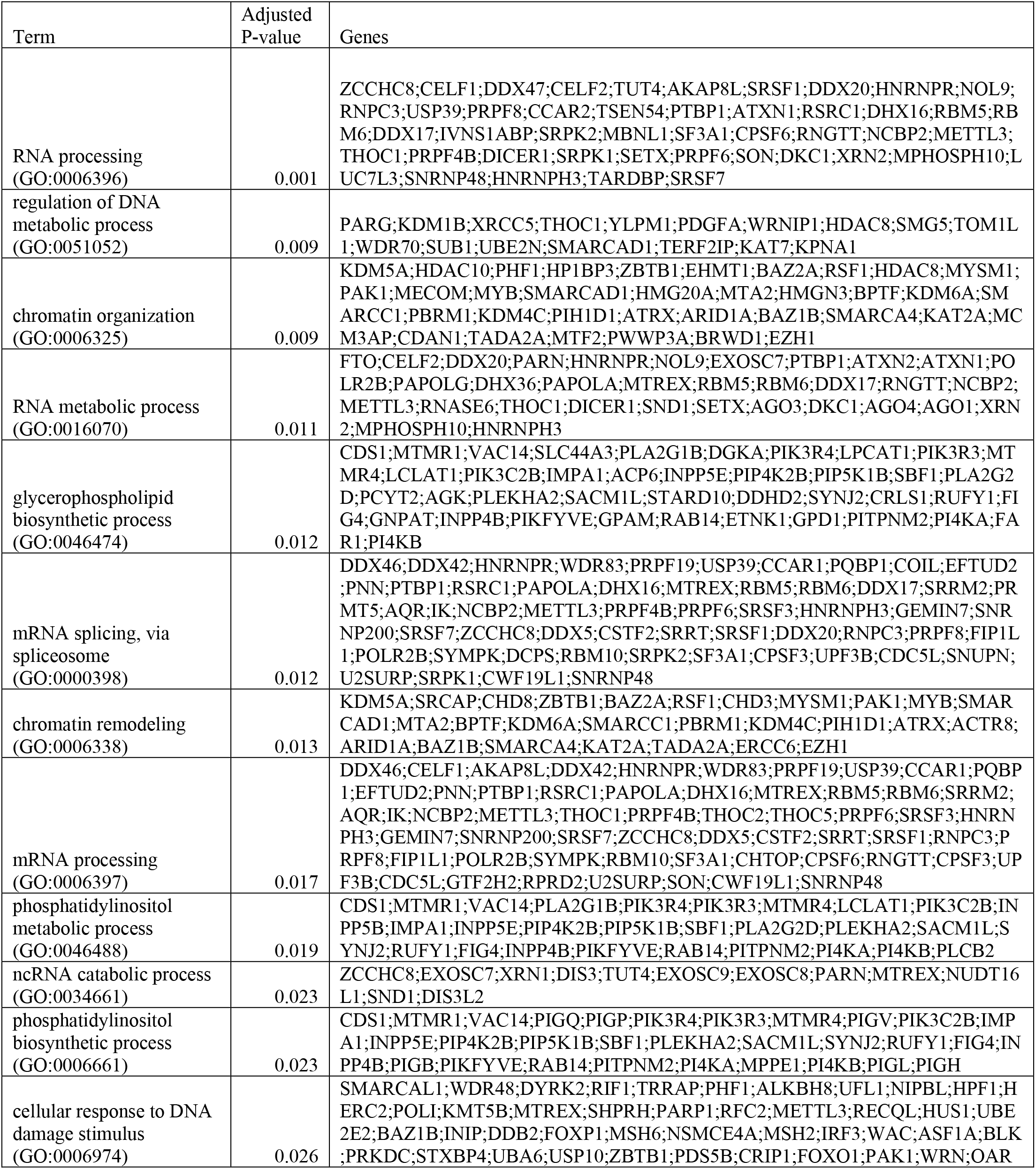
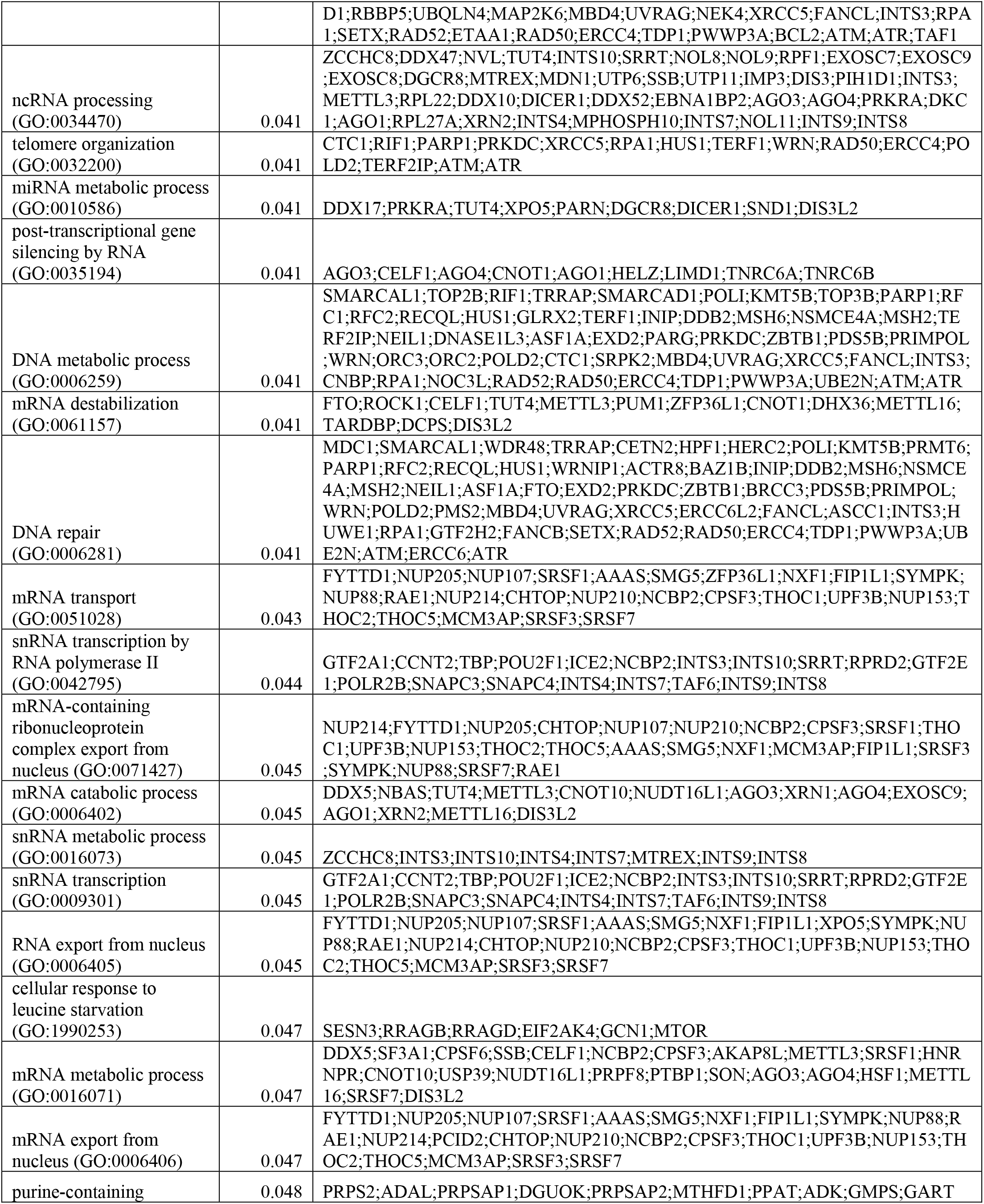
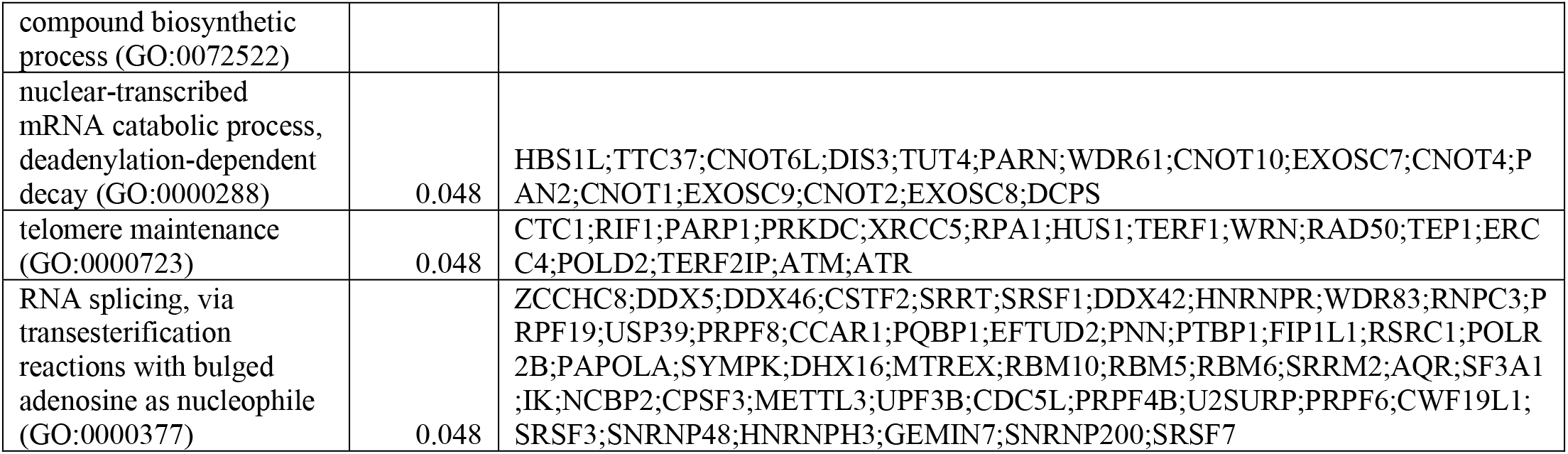
Enrichr analysis resulting from the genes selected for the Asymptomatic vs Progressor classification. Adjusted *p*-value <0.05 are displayed.

## SUPPLEMENTARY METHODS

### Preprocessing of H&E images

Prior to model development, images were preprocessed as described in [24] to detect image foreground by thresholding the red channel with a value of 230; filling holes; and removing artifacts by applying an image erosion and dilation with a disk structuring element with a radius of 3. Then, image patches sized 32×32 pixels at 20x magnification were passed through ResNet50 pretrained on ImageNet until the 2nd residual block. The resultant 8×8×256 feature block was collapsed into a 1×256 feature vector (later called embedded instances) by averaging along the first two dimensions.

### Attention-based multiple instance learning

Attention-based automatically learns to weight embedded instances into a bag-level feature vector that can be subsequently classified. In a single image, we sampled patches (instances) from the lung section image and performed automated feature extraction to generate the instance embeddings. Each embedded instance was weighted, summed, and combined into a single, image-level embedding to allow classification.

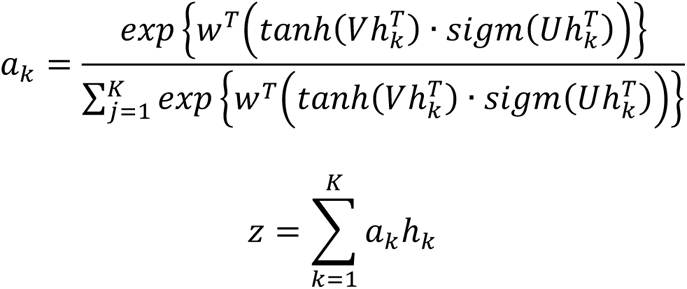

The attention mechanism implementation consists of a simple two-layer fully connected network which passes each instance embedding (hk) though two parallel layers of the network (V,U), applies tanh and sigmoid activation functions to respective outputs, dots the two results, then passes the activation though the second layer (wT), which maps the vector into a single value, its attention weight (ak). The weighted sum of each embedded instance and its attention weight yields a bag-level instance (z). The parameters (V,U,w) for this for this neural network are automatically learned through training of the model.

In addition to performing better than instance-based and embedding-based max and mean pooling approaches, the resulting instance weights allow the model to be interpretable in that the relative magnitudes of instance weights directly correspond to the instance’s relative contribution to the overall classification of the bag.

### Further details regarding the classifiers used with lung cytokines, chemokines, and IgG antibodies

We have used one linear classifier, l1-regularized logistic regression, and one non-linear classifier XGBoost. Logistic regression model parametrizes the conditional distribution of the class labels using a linear combination of the input features, *x* ∈ *R^p^*, i.e. *P*(*Y* = 1|*X* = *x*, *β*, *b*) = (1 + exp(−*β^T^x* − *b*))^-1^. Let *x*^(*i*)^ denote the input features and *y*^(*i*)^ ∈ {−1,1} the corresponding label of the i^th^ sample. Given N independent and identically distributed samples {*x*^(*i*)^, *y*^(*i*)^}*^N^_i=_*_1_ and the regularization amount *λ*, to estimate the parameters the sum of the negative log-likelihood and *ℓ*_1_ norm of the weights is minimized i.e. ∑ log(1 + *e*^*-y*(*i*)(*βTx*(*i*)+*b*)^) + *λ*‖*β*‖_1_. XGBoost like gradient tree boosting, [61] is a method for iteratively fitting a set of decision trees whose additive combination determines the predicted class probabilities. At each iteration, the first and second derivatives of the resulting loss function from the existing set of trees guide the selection of the new tree [33]. This process is repeated till the desired number of trees is reached.

### Comparison between Mann–Whitney U-statistic and Welch’s t-test

In this work, motivated by identifying diagnostically powerful genes, we have preferred AUC analysis of the gene expression profiles to a more standard approach such as the t-test. To measure the sensitivity of the selected genes to the statistical test used we have contrasted the selections of the one-sided Mann–Whitney U-statistic with those of one-sided Welch’s t-test. For the comparison between Asymptomatic and all remaining mice (Controllers, Progressors, and non-infected controls) within the Experiment 1, one-sided Welch’s t-test identified 1133 genes that are expressed at significantly (FDR q < 0.05) as opposed to 105 genes identified using the one-sided Mann–Whitney U-statistic. The resulting 1133 selections from Welch’s t-test contained all 105 selections resulting from the Mann–Whitney U-statistic. However, using the Welch’s t-test resulted in genes with diagnostically less powerful to be selected. The gene with the least diagnostic potential had 0.630 AUC in the set resulting from Welch’s t-test as opposed to 0.844 AUC in the set resulting from Mann–Whitney U-statistic (Supplementary File 1). When 1133 selections from Welch’s t-test input to Enrich, 41 statistically significant (adjusted *p* < 0.05) GO terms were identified and only two of them overlapped with 11 GO terms identified with Mann–Whitney U-statistic namely, “regulation of B cell receptor signaling pathway” and “regulation of antigen receptor-mediated signaling pathway” (Supplementary File 1). Even if selections from theWelch’s t-test contain the previous 105 selections, the order of magnitude higher number of input genes to Enrichr might have decreased the statistical significance of the previously identified pathways.

